# Interferon-α promotes neo-antigen formation and preferential HLA-B-restricted antigen presentation in pancreatic β-cells

**DOI:** 10.1101/2023.09.15.557918

**Authors:** Alexia Carré, Zhicheng Zhou, Javier Perez-Hernandez, Fatoumata Samassa, Christiana Lekka, Anthony Manganaro, Masaya Oshima, Hanqing Liao, Robert Parker, Annalisa Nicastri, Barbara Brandao, Maikel L. Colli, Decio L. Eizirik, Marcus Göransson, Orlando Burgos Morales, Amanda Anderson, Laurie Landry, Farah Kobaisi, Raphael Scharfmann, Lorella Marselli, Piero Marchetti, Sylvaine You, Maki Nakayama, Sine R. Hadrup, Sally C. Kent, Sarah J. Richardson, Nicola Ternette, Roberto Mallone

## Abstract

Interferon (IFN)-α is the earliest cytokine signature observed in individuals at risk for type 1 diabetes (T1D), but its effect on the repertoire of HLA Class I (HLA-I)-bound peptides presented by pancreatic β-cells is unknown. Using immunopeptidomics, we characterized the peptide/HLA-I presentation in *in-vitro* resting and IFN-α-exposed β-cells. IFN-α increased HLA-I expression and peptide presentation, including neo-sequences derived from alternative mRNA splicing, post-translational modifications - notably glutathionylation - and protein *cis*-splicing. This antigenic landscape relied on processing by both the constitutive and immune proteasome. The resting β-cell immunopeptidome was dominated by HLA-A-restricted ligands. However, IFN-α only marginally upregulated HLA-A and largely favored HLA-B, translating into a major increase in HLA-B-restricted peptides and into an increased activation of HLA-B-restricted vs. HLA-A-restricted CD8^+^ T-cells. A preferential HLA-B hyper-expression was also observed in the islets of T1D vs. non-diabetic donors, and we identified islet-infiltrating CD8^+^ T-cells from T1D donors reactive to HLA-B-restricted granule peptides. Thus, the inflammatory milieu of insulitis may skew the autoimmune response toward epitopes presented by HLA-B, hence recruiting a distinct T-cell repertoire that may be relevant to T1D pathogenesis.

## Introduction

Type I interferons (IFNs), notably IFN-α, play a crucial role in anti-viral immune responses by driving infected cells to express IFN-stimulated genes, thus limiting viral replication and spreading^1^. Type I IFNs are also central in modulating innate immune responses and activating adaptive immunity^1^. They boost B-cell differentiation and antibody (Ab) production^2^, mature antigen-presenting cells^2^, and upregulate their expression of human leukocyte antigen (HLA) Class I (HLA-I) and Class II along with costimulatory molecules^3^. Upon detection of microbial components, macrophages and dendritic cells (DCs; mainly plasmacytoid DCs) secrete IFN-α, leading to additional cytokine and chemokine release. IFN-α also increases HLA-I expression directly on infected cells^4^, thus augmenting their antigenic visibility, and enhance the cytolytic activity and survival of natural killer (NK) and CD8^+^ T-cells^5–7^.

Besides these potent anti-viral effects, IFNs can also promote autoimmunity^8^. In type 1 diabetes (T1D), transcriptomics investigations on the peripheral blood of children genetically at risk for disease revealed a strong type I IFN signature^9^, preceding seroconversion^10,11^. At the tissue level, IFN-stimulated genes are found overexpressed only in infiltrated islets of recent-onset T1D patients^12^, in line with an enhanced pancreatic IFN-α expression^13,14^. These IFN signature may reflect exposure to viral infections, with Coxsackieviruses B proposed as possible environmental triggers of islet autoimmunity^15^, or to other “danger signals”.

IFN-α^13^ and IFN-stimulated gene expression^16,17^ colocalize with HLA-I hyper-expression, which is a histopathological hallmark of T1D in insulin (INS)-containing islets^18^. More recently, a causal relationship between IFN-α, endoplasmic reticulum (ER) stress, and HLA-I upregulation has been demonstrated in β-cells^4,19^. The increased surface HLA-I expression on β-cells may be a driver of T1D pathogenesis, as self-reactive CD8^+^ T-cells must recognize peptide-HLA-I (pHLA-I) complexes on β-cells to trigger lysis. Additionally, both IFN-α and ER stress may impact the repertoire of HLA-I-presented peptides (so called immunopeptidome), as is the case for IFN-γ^20–22^. Indeed, IFN-α promotes the generation of mRNA splice variants^19,23^, which can result in the presentation of neo-epitopes spanning novel splicing sites. This loss of immune ignorance may also rely on other cytokine-induced pathways of neo-epitope formation^20,24^, e.g. post-translation modifications (PTMs), alternative transcription start sites generating an INS defective ribosomal product (DRiP)^25^ and peptide *cis*-splicing, i.e. the fusion of non-contiguous protein fragments.

IFN-α may thus modulate the antigenic cargo offered to T-cells. Against this background, elucidating how IFN-α shapes the immunopeptidome of β-cells is crucial to further understand the antigenic drivers of T1D. We have therefore used immunopeptidomics strategies applied to human β-cells treated or not with IFN-α to define the antigenic display driving autoimmune T-cell recognition.

## Results

### ECN90 β-cells exposed to IFN-α increase surface HLA-I expression and peptide presentation

To understand changes in the profile of HLA-I-presented peptides in β-cells exposed to IFN-α, we cultured the ECN90 β-cell line^26^ with or without IFN-α for 18 h. pHLA-I complexes were immunoprecipitated from cell lysates, and peptides identified by liquid chromatography tandem mass spectrometry (LC-MS/MS). ECN90 β-cells exposed to IFN-α yielded more peptides than untreated cells (Fig. 1A-B), with respectively 91.1% and 74.9% sequences in the expected 8-14 amino-acid (aa) range for HLA-I ligands (Fig. 1A, E). This observation reflected an IFN-α-induced upregulation of surface HLA-I (Fig. 1C-D). Unsupervised Gibbs clustering uncovered the peptide motifs of HLA-I alleles expressed by ECN90 β-cells (Supplementary Fig. 1). The predominant group 1 identified the HLA-A*03:01 (HLA-A3 from hereon) motif, group 2 corresponded to a mixture of HLA-A*02:01 (HLA-A2), -C*03:04 (-C3) and -C*07:01 (-C7) motifs, and group 3 matched the related motifs of HLA-B*40:01 (-B40) and -B*49:01 (-B49). Peptides were identified and filtered by a multi-step bioinformatics pipeline (Fig. 1E). The reference database used for the identification of genome-templated sequences was preliminarily complemented with translated mRNA isoforms enriched in human islets exposed or not to IFN-α^19^ and generating aa neo-sequences; and with *INS* open-reading frames covering published neo-epitopes^25^. Following selection for 8-14 aa length, and identification of mRNA variants, all sequences underwent filtering based on enriched expression of source proteins in β-cells. Peptides carrying PTMs were analyzed separately. All those spectra whose peptide interpretation did not match genome-templated, RNA-spliced or PTM sequences were searched for putative proteasomal *cis*-spliced peptides using the MARS algorithm^27^. In total, this workflow led to the identification of 4 mRNA splice candidates (<u>Supplementary Data 1</u>), 784 conventional peptides (<u>Supplementary Data 2</u>), 247 sequences carrying PTMs (<u>Supplementary</u> <u>Data 3</u>), 11 *cis*-spliced sequences (<u>Supplementary Data 4</u>). HLA-I restrictions were then predicted using NetMHCpan 4.1a^28^. For all peptide categories, more HLA-I binders were identified in cells exposed to IFN-α (Fig. 1E), in line with the induced surface HLA-I upregulation.

**Figure 1.**
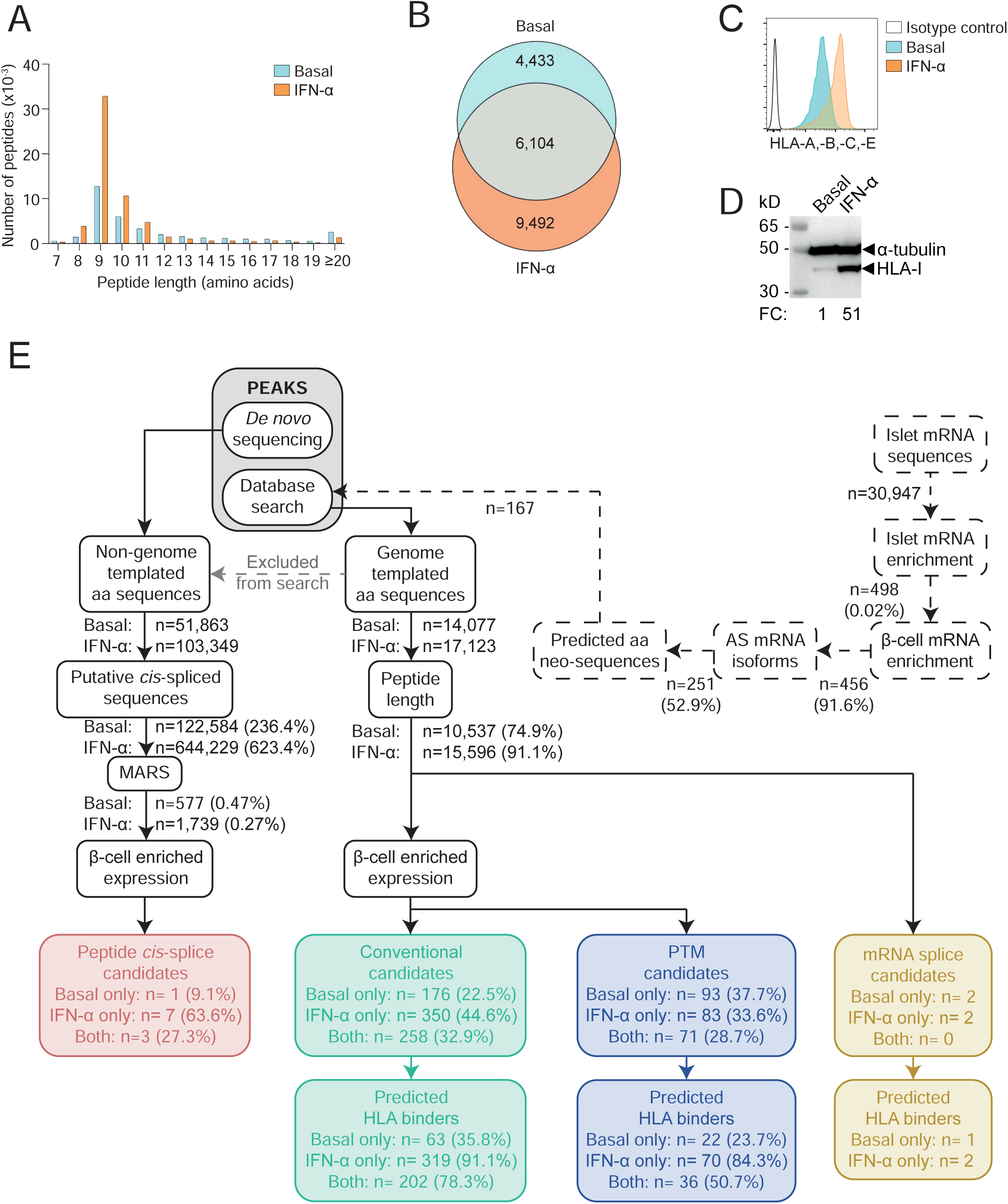
Immunopeptidome profiling of HLA-I-bound peptides presented by ECN90 β-cells. **A.** Length distribution of HLA-I-eluted peptides from ECN90 β-cells under basal (blue) and IFN-α-stimulated conditions (orange). Four biological replicates (3.5-4.5×10^8^ cells/each) were acquired for each condition, and unique peptides across replicates were counted. **B.** Number of 8-14mer peptides eluted from the above conditions. **C.** HLA-I expression detected by flow cytometry on ECN90 β-cells in basal (blue) and IFN-α-stimulated conditions (orange) using W6/32 Ab. **D.** HLA-I heavy chain expression detected with HC10 Ab by Western blot in whole cell lysates of ECN90 cells exposed or not to IFN-α, with α-tubulin bands as loading controls and normalized HLA-I fold change (FC) values indicated. **E.** Bioinformatics analysis pipeline. Predicted aa neo-sequences from RNAseq datasets of human islets (dashed lines) were appended to the database used for immunopeptidome search. Database-matched sequences identified by PEAKS (grey box) were sequentially filtered based on their length, on whether they matched mRNA variants (yellow boxes; peptides listed in <u>Supplementary Data 1</u>) and on the enriched expression of their source proteins in β-cells. Conventional candidates (green boxes; <u>Supplementary Data 2</u>) and sequences carrying PTMs (violet boxes; <u>Supplementary</u> <u>Data 3</u>) were separated. HLA-I-binding predictions were performed using NetMHCpan4.1a (peptide motifs detailed in Supplementary Fig. 1). In parallel, non-genome-templated peptides were interpreted as potential *cis*-spliced candidates and fed into the MARS algorithm, followed by filtering according to the enrichment of their source proteins in β-cells (red box, <u>Supplementary Data 4</u>).

Collectively, these results show that IFN-α increases surface HLA-I expression and peptide presentation.

### The immunopeptidome of ECN90 β-cells is dominated by insulin granule proteins and its diversity is increased by IFN-α

Next, we investigated the source proteins of the β-cell-enriched peptides identified. IFN-α treatment increased not only the number of peptides presented, but also the diversity of source proteins, with 35.6% (58/163) of them appearing only in IFN-α-treated cells as compared to 6.1% (10/163) specific to untreated cells (Supplementary Fig. 2A). Accordingly, 19 source proteins in IFN-α-treated cells yielded ≥2-fold more peptides (fold change log_2_FC>1) than in the control condition (Fig. 2A, Supplementary Fig. 2A). Conversely, only 8 source proteins displayed a similar log_2_FC<1 in untreated vs. IFN-α-treated cells. In both conditions, the 5 source proteins yielding more peptides are known to either localize in secretory granules, i.e. chromogranin A (CHGA, n=132), INS (n=64), secretogranin 5 (SCG5, n=34); or to be associated with granule transport, i.e. kinesin-like protein (KIF1A, n=34) and microtubule-associated protein 1B (MAP1B, n=26). Conversely, some known β-cell antigens were underrepresented (Supplementary Fig. 2A), i.e. zinc transporter 8 (SLC30A8, n=5), glutamate decarboxylase (GAD2, n=3) and islet amyloid polypeptide (IAPP, n=1), or completely absent, i.e. islet-specific glucose-6-phosphatase catalytic subunit-related protein (IGRP) and INS DRiP^25^. Novel granule proteins were also identified, i.e. chromogranin B (CHGB, n=6), secretogranin 2 and 3 (SCG2, n=11; SCG3, n=10).

**Figure 2.**
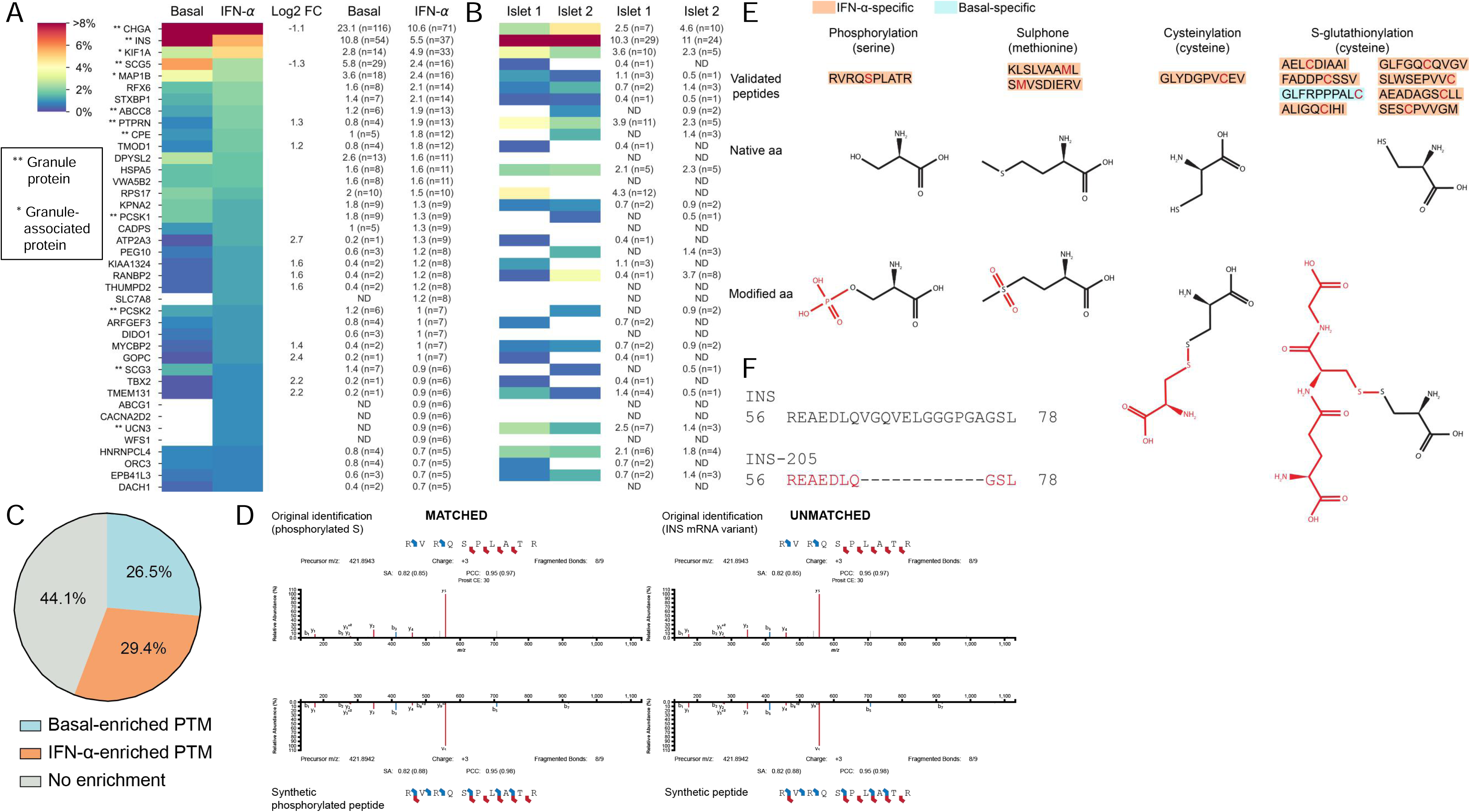
Immunopeptidome of ECN90 β-cells exposed or not to IFN-α and validation of HLA-I-eluted candidate neo-epitopes. **A-B**. Heatmap of the relative representation of the top 40 source proteins in ECN90 β-cells (A) and human islets (B), ranked according to the number of peptides detected in the IFN-α-treated condition. The color scale is proportional to the number of peptides identified for each protein out of the total number of peptides in a given condition, expressed as percentage. Only conventional and PTM sequences are included. Peptides carrying PTMs were counted as such only for PTMs defined as likely biological (they were otherwise counted as unmodified); *cis*-spliced peptides were excluded. Percent values and peptide numbers are listed on the right. Source proteins enriched in IFN-α-treated cells (log2 fold change, FC ≥ 1) or in basal condition (log2 FC ≤ 1) are indicated for ECN90 β-cells. The complete heatmap is provided in Supplementary Fig. 2. HLA-I-bound peptides eluted from primary human islets are listed in <u>Supplementary Data 5</u> and compared with those eluted from ECN90 β-cells in <u>Supplementary Data 6</u>. **C.** Global PTM enrichment in basal and IFN-α-treated condition. A detailed list is provided in <u>Supplementary Data 7</u>. **D.** Spectral matching examples of matched (left) and unmatched (right) synthetic peptides compared to the initial peptide identification using an online tool (https://www.proteomicsdb.org/use). **E.** Validated post-translationally modified peptides and representation of the native and modified aa. The PTM is indicated in red. A detailed list is provided in <u>Supplementary Data 3</u>. **F.** Peptide alignment of INS and INS-205 mRNA variant. The sequence of the HLA-eluted variant is indicated in red and correspond to a sequence spliced out as compared to the canonical INS mRNA.

To validate the results obtained on ECN90 β-cells, HLA-I-bound peptides were identified from two untreated HLA-A2/A3^+^ primary human islet preparations from non-diabetic donors. Despite the limited (∼100-fold lower) amount of starting material and the dilution of β-cell-derived peptides with those from other endocrine cells, 342 β-cell-enriched conventional peptides were retrieved from these islet preparations (<u>Supplementary Data 5</u>), i.e. only 2-fold less than in ECN90 β-cells (n=784). Across the 2 islet samples, the source proteins (n=72) of the peptides retrieved overlapped by 44% with those from ECN90 β-cells (n=163) (Supplementary Fig. 2B). Interestingly, even though glucagon (GCG)-secreting α-cells account for only half the number of β-cells in human islets^29^, the most represented endocrine-enriched protein was GCG (n=26 and 30 peptides from islet sample 1 and 2, respectively; n=43 unique peptides; Supplementary Fig. 2C). The other dominant source proteins were otherwise the same as in ECN90 β-cells, notably CHGA (n=7+10, 12 unique peptides) and INS (n=29+24, 33 unique peptides; Fig. 2B, <u>Supplementary Data 5</u>). In total, 40/117 (34%) HLA-A2/-A3-restricted peptides identified in primary islets were also found in ECN90 β-cells (<u>Supplementary Data 6</u>), which was only slightly lower than the overlap between the two islet preparations (56/117 shared peptides, 48%).

Collectively, these results show that IFN-α increases not only the number of peptides displayed by HLA-I, but also their diversity in terms of source proteins. Granule-contained and granule-associated proteins are abundantly represented, with some novel ones (CHGB, SCG2, SCG3) identified.

### The immunopeptidome of ECN90 β-cells includes PTM, mRNA splicing and peptide *cis*-splicing neo-sequences and is shaped by both the constitutive and immuno-proteasome

We subsequently focused on candidate neo-epitopes generated by PTMs and mRNA or peptide *cis*-splicing. Thirty-four PTMs were identified on HLA-I-eluted ligands from ECN90 β-cells (<u>Supplementary Data 7</u>). Several of them (15/34, 44.1%) were found in the same proportion in both basal and IFN-α-treated samples (Fig. 2C). Since most PTM neo-epitopes are generated under cell stress conditions^30^, we focused on those specifically enriched after IFN-α treatment (10/34, 29.4%). In order to not dismiss the opposite scenario of physiological PTMs that may be lost in an inflammatory environment, we also included those enriched under basal conditions (9/34, 26.5%). The second selection criterion was the possibility to chemically introduce the modification on synthetic peptides for MS validation, excluding PTMs that naturally arise during peptide synthesis or MS acquisition, e.g. oxidations and all PTMs found at the more vulnerable N- and C-terminal positions. Following these criteria, we retained for further study lysine acetylation, cysteinylation, deamidation, glutathione disulfide (S-glutathionylation), phosphorylation and sulphone; and synthesized predicted HLA-A and -B binders in both modified and unmodified form. We then compared the spectrum of the identified PTM peptides: a) with that of the synthetic PTM peptides; and b) with the spectrum of the corresponding peptides synthesized in their native form (Fig. 2D). In this second case, a spectral match would indicate an artifactual modification that is experimentally introduced. Only 12/31 *bona fine* biological PTM peptides were thus validated, carrying: phosphorylation (n=1 peptide), sulphone (n=2), cysteinylation (n=1) and S-glutathionylation (n=8) (Fig. 2E, <u>Supplementary</u> <u>Data 3</u> and <u>Supplementary Data 7</u>). Interestingly, 11/12 PTM peptides were exclusively found in IFN-α-treated cells. The immunopeptidome of human islets did not retrieve glutathionylated peptides.

Spectral matching of the 3 HLA-I-restricted mRNA splice candidates identified validated only one HLA-B40/B49-restricted INS-205 sequence (Fig. 2F), which was exclusively found in IFN-α-treated cells (<u>Supplementary Data 1</u>). We further assigned 11 peptides which could hypothetically be explained by *cis*-splicing, either at the RNA or protein level. We could validate most spectra by matching to spectra of the synthetic counterpart (9/11, 81.8%), with some isoleucine/leucine ambiguities not resolvable by MS (<u>Supplementary Data 4</u>). Altogether, 12 PTM peptides, 1 mRNA splice and 9 candidate *cis*-spliced peptides were validated by spectral matching.

To explore whether the generation of the identified peptides was proteasome-dependent, we characterized changes in the immunopeptidome of ECN90 β-cells exposed or not to IFN-α and (immuno)proteasome inhibitors. The catalytic subunits of the constitutive and immuno-proteasome were found expressed by ECN90 β-cells, with the latter upregulated by IFN-α treatment as expected (Supplementary Fig. 3A). Treatment with carfilzomib, an inhibitor of the β5 proteasome subunit, or with ONX-0914, an inhibitor of the β5i immunoproteasome subunit with minimal cross-reactivity for its conventional β5 counterpart, led to a significantly decreased chymotrypsin-like activity (Supplementary Fig. 3B). The HLA-I-eluted ligands were filtered and processed following the previous bioinformatics pipeline. The β-cell-enriched peptides retrieved across conditions clustered into distinct subgroups (Supplementary Fig. 3C, <u>Supplementary Data 8</u>). Some peptides (cluster 6) were specific to untreated β-cells and remained unaffected by constitutive proteasome inhibition. Conversely, peptides in cluster 3, also specific to untreated β-cells, were proteasome-dependent. Peptides in clusters 9 and 10 were instead largely specific to IFN-α-treated β-cells and, respectively, dependent and independent on immuno-proteasome catalysis. Of note, cluster 10 can visually be divided in peptides dependent (bottom half) or not (top half) on constitutive proteasome. Peptides in cluster 2 were dependent on constitutive or immuno-proteasome, regardless of IFN-α exposure. Similarly, the presentation of certain peptides was only enriched in β-cells treated with carfilzomib, regardless of IFN-α exposure (cluster 1). Peptides enriched upon immunoproteasome inhibition were also observed (cluster 8). Cluster 7, 4 and 5 were the most surprising. Cluster 7 depended on the exposure to either carfilzomib alone or a combination of IFN-α and carfilzomib/ ONX-0914. Cluster 4 was enriched in β-cells either untreated or exposed to both IFN-α and ONX-0914, while cluster 5 was enriched in basal conditions, regardless of carfilzomib exposure, or β-cells treated with both IFN-α and ONX-0914. Of note, the preproINS (PPI)_15-24_ peptide was not affected by (immuno)proteasome inhibition and was consistently identified in all conditions.

Collectively, these results show that the immunopeptidome is shaped by both the constitutive and immuno-proteasome expressed by β-cells.

### IFN-α skews peptide presentation toward HLA-B ligands

Another distinctive feature of IFN-α-treated ECN90 β-cells was the significant enrichment in HLA-B binders. In untreated cells, only 8% (41/502) of peptides were predicted HLA-B binders (-B40, -B49 or both), which increased to 34% (226/672) in IFN-α-treated cells (Fig. 3A; detailed in Supplementary Fig. 4). A similar but lesser effect was noted for HLA-C [5% (25/502) vs. 8% (56/672)], while the number of HLA-A and HLA-E ligands remained stable [41% (208/502) vs. 41% (277/672) and 0.8% (4/502) vs. 1.0% (7/672), respectively]. Thus, HLA-B ligands were the main contributor to the enriched peptide display induced by IFN-α.

**Figure 3.**
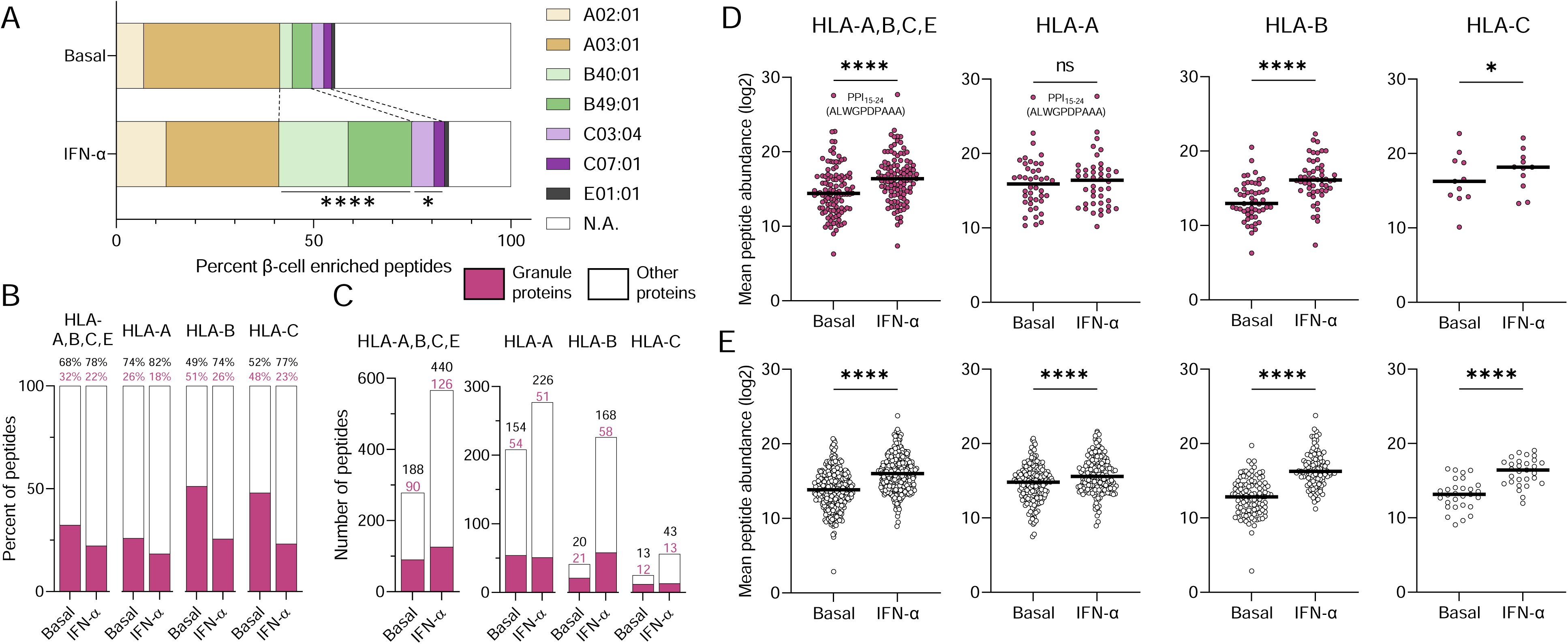
HLA-I restrictions of the immunopeptidome of ECN90 β-cells exposed or not to IFN-α. **A.** Relative distribution of predicted HLA-I ligands for each allele expressed by ECN90 β-cells in basal and IFN-α-treated conditions. \*\*\*\**p*<0.0001 and \**p*=0.027 by Fisher exact test. Predicted HLA-E*01:01-restricted peptides are listed in <u>Supplementary Data 9</u>. **B-C.** Percent proportion (B) and number of peptides (C) originating from granule-contained and other proteins in basal and IFN-α-treated conditions. A heatmap of the source proteins of HLA-A- and HLA-B-restricted peptides is provided in Supplementary Fig. 4. **D-E.** Average total abundance of conventional peptides originating from granule-contained (D) and other proteins (E). The peptides were identified by PEAKS and quantified by Progenesis. The PPI_15-24_ sequence of the most abundant peptide identified in both conditions is indicated. Horizontal bars represent median values. \*\*\*\**p*<0.0001 and \**p*<0.05 by Wilcoxon test.

Granule proteins contributed 32% of the HLA-I immunopeptidome in untreated ECN90 β-cells, which decreased to 22% upon IFN-α treatment (Fig. 3B), reflecting a dilutional effect due to the higher increase of peptides from other sources (Fig. 3C). Nonetheless, when analyzing HLA-A, -B and -C ligands separately, IFN-α increased the number of peptides from granule proteins only for HLA-B (Fig. 3C). The IFN-α-driven diversification of the immunopeptidome was also larger for HLA-B, which yielded 5.5-fold more peptides (226 vs. 41 in IFN-α-treated vs. untreated cells) compared to 1.3-fold (277 vs. 208) for HLA-A; and 4.2-fold (98 vs. 23) more source proteins compared to 1.2-fold (107 vs. 89) for HLA-A ligands (Fig. 3C, Supplementary Fig. 4). In terms of peptide abundance (Fig. 3D-E), the increase was also more evident for HLA-B ligands, particularly for granule-derived peptides, which did not significantly increase for HLA-A. The previously described HLA-A2-restricted preproINS (PPI)_15-24_ peptide^20^ was the most abundantly presented, and was not affected by IFN-α treatment.

Using stringent criteria (i.e. no alternative restriction predicted), 7 conventional peptides predicted to bind HLA-E*01:01 were also identified, none of which was found exclusively in the basal condition, in line with the reported HLA-E upregulation induced by IFN-α^19^. A more extensive list of putative HLA-E ligands is provided in <u>Supplementary Data 9</u>.

Collectively, IFN-α skews antigen presentation toward HLA-B-restricted peptides.

### IFN-α preferentially upregulates HLA-B expression without inducing dedifferentiation or reducing proINS synthesis

Next, we considered whether the increased presentation of HLA-B ligands induced by IFN-α could reflect preferential HLA-B upregulation. Indeed, *HLA-B* gene expression increased 42-fold and 226-fold in IFN-α-treated and IFN-γ-treated ECN90 β-cells, respectively, while *HLA-A* was only marginally upregulated and *HLA-C* displayed an intermediate increase (Fig. 4A). We then validated HLA-A, -B and -C Abs for their staining specificity, using HLA-I^−^ K562 cells transduced with different HLA-I alleles. Using flow cytometry (Supplementary Fig. 5A), HLA-A Ab ARC0588 and HLA-C Ab DT-9 only recognized their respective alleles, while HLA-B Ab JOAN-1 displayed some cross-reactivity with HLA-A alleles of low prevalence and a lesser reactivity to HLA-C variants. The HLA-A Ab ARC0588 was also functional and specific by Western blot (Supplementary Fig. 5B), while the JOAN-1 and DT-9 Abs did not detect any band. Another Ab HC10, which was not functional by flow cytometry, preferentially recognized HLA-B (Supplementary Fig. 5B), with minimal cross-reactivity for HLA-C. Using these validated Abs, protein expression confirmed that, while only HLA-A was expressed under basal conditions, both IFN-α and IFN-γ preferentially upregulated HLA-B and, to a minor extent, HLA-C, on ECN90 β-cells (Fig. 4B-C).

**Figure 4.**
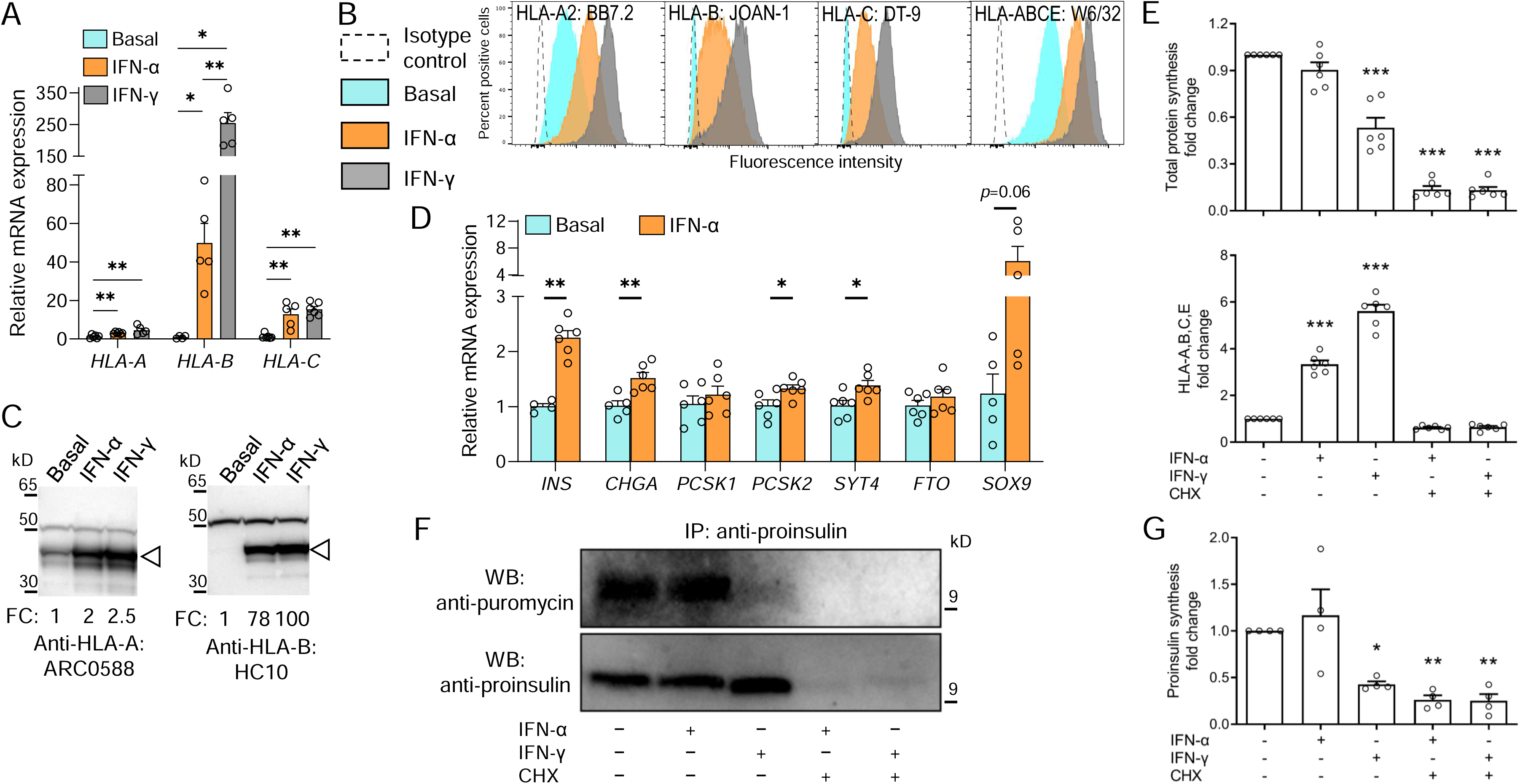
IFN-α preferentially upregulates HLA-B expression without inducing dedifferentiation or reducing proINS synthesis. **A.** Relative mRNA expression of *HLA-A*, *HLA-B* and *HLA-C* alleles in ECN90 β-cells exposed or not to IFN-α or IFN-γ for 24 h. *GAPDH* was used as an internal normalizing control, and each gene was normalized to the basal sample. Data represent mean±SEM of 5 biological replicates. \*\**p*<0.01 and \**p*<0.05 by Mann-Whitney U test. **B-C.** Protein expression of HLA-A, -B and -C in in ECN90 β-cells exposed or not to IFN-α or IFN-γ, as detected by surface flow cytometry (B) and Western blot (C) using the indicated Abs validated for their specificity (see Supplementary Fig. 5). For Western blotting, the arrowhead indicates the HLA-I heavy chain band, with the top band indicating the α-tubulin loading control and normalized HLA-I fold change (FC) values indicated. **D.** Relative mRNA expression of β-cell identity genes in ECN90 β-cells exposed or not to IFN-α. *PPIA* was used as internal normalizing control, and data representation is the same as in panel A. \*\**p*<0.01 and \**p*<0.05 by Mann-Whitney U test. **E.** Puromycin incorporation in newly synthesized total proteins (top) and HLA-A/B/C/E expression (bottom), detected by flow cytometry and expressed as fold change compared to the basal sample (first column). Inhibition of protein synthesis by cycloheximide (CHX) provided negative controls (last 2 columns). Data represent mean±SEM of 6 biological replicates. \*\*\**p*<0.0001 by one-way ANOVA. **F.** Puromycin incorporation in newly synthesized proINS, detected by proINS immunoprecipitation followed by Western blot for puromycin (top; corresponding to newly synthesized proINS) and proINS (bottom; corresponding to total proINS). A representative experiment out of 4 performed is shown. **G.** Puromycin incorporation in newly synthesized proINS, detected by Western blot and expressed as fold change compared to the basal sample (first column). Data represent mean±SEM of 4 biological replicates. \**p*<0.03 and \*\**p*<0.002 by one-way ANOVA.

We then analyzed the effect of IFN-α on the expression of β-cell identity genes (Fig. 4D). Despite a variable upregulation of the progenitor marker *SOX9*, several β-cell identity markers were also more consistently upregulated by IFN-α, i.e. *INS*, *CHGA*, *PCSK2*, *SYT4*, suggesting that de-differentiation was not induced. Using puromycin treatment (which is incorporated during protein synthesis) and an anti-puromycin Ab to detect newly synthesized proteins^31^, IFN-α did not downregulate overall *de-novo* protein synthesis (Fig. 4E), while it upregulated HLA-I translation, as expected. *De-novo* proINS synthesis was also unaffected (Fig. 4F-G), at variance with the downregulating effect of IFN-γ on both total protein and proINS synthesis (Fig. 4E-F-G).

Collectively, these results document a preferential HLA-B gene and protein upregulation by IFN-α and, to a larger extent, IFN-γ. IFN-α neither induced β-cell de-differentiation nor downregulated proINS expression.

### Preferential HLA-B hyper-expression in the islets of T1D donors

These observations prompted us to test whether the histopathological hallmark of HLA-I hyper-expression in the islets of T1D patients^18^ could also preferentially involve HLA-B. To this end, we stained pancreas tissue sections of T1D (n=6) and non-diabetic donors (n=4) from the Network for Pancreatic Organ Donors with Diabetes (nPOD) and Exeter Archival Diabetes Biobank (EADB; Supplementary Table 1), using the previously validated HLA-A Ab ARC0588, HLA-B Ab HC10 (the other HLA-B Ab JOAN-1 was not functional by immunofluorescence) and the HLA-A/B/C/E Ab EMR8-5^18^ (Fig. 5A-B-C). As expected, an increased HLA-A/B/C/E expression was detected in INS-containing islets (ICIs) from T1D compared to non-diabetic (ND) donors, both in β-cells and α-cells (Fig. 5D). However, HLA-B was upregulated to a greater extent than HLA-A. Conversely, HLA-A/B/C/E, HLA-A and HLA-B expression in α-cells from T1D islets devoid of β-cells (INS-deficient islets, IDIs; Supplementary Fig. 6A) was lower than in α-cells from T1D ICIs, and comparable to that of ND ICIs (Fig. 5D). Moreover, assessment of individual β- and α-cell HLA-I expression in T1D donors with increasing disease duration showed that HLA-B expression was highest in 4 of the 5 donors (Supplementary Fig. 6B-C) and, as for other HLA-Is, was sustained in ICIs, but reduced in IDIs with increasing disease duration (Supplementary Fig. 6D).

**Figure 5.**
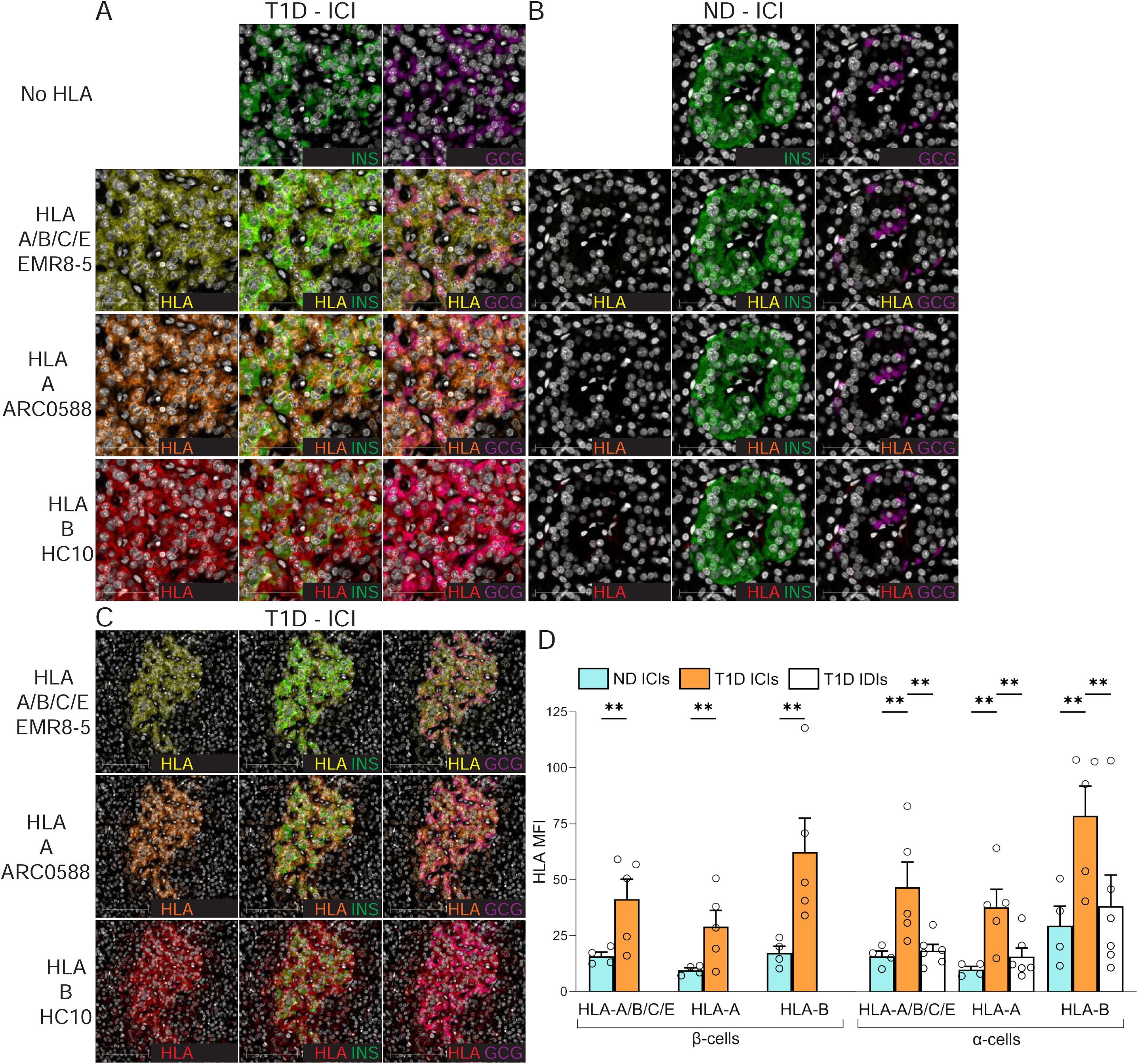
HLA-B vs. HLA-A hyper-expression in the islets of T1D and non-diabetic (ND) cases. **A-B.** Representative immunofluorescence images of ICIs from T1D case nPOD 6396 (A) and ND case nPOD 6160 (B; all cases listed in Supplementary Table 1), stained with DAPI only (first row) or for HLA-A/B/C/E (yellow; second row), HLA-A (orange; third row) and HLA-B (red, fourth row), alone (first column) or in combination with INS (green, second column) or GCG (violet, third column). Scale bar 50 µm. **C.** Whole ICI images from the same T1D case nPOD 6396, scale bar 100 µm. **D.** Immunofluorescence quantification of HLA-I mean fluorescence intensity (MFI) for HLA-A/B/C/E, HLA-A and HLA-B in β-cells (left) and α-cells (right) from ICIs of ND (blue; n=4) and T1D cases (orange; n=5); and in α-cells from IDIs of T1D cases (white; n=6). Bars represent mean+SEM values. \*\**p*≤0.005 by 2-way ANOVA. T1D IDI images and individual quantifications for each T1D donor are provided in Supplementary Fig. 6.

Collectively, these results confirm the HLA-I hyper-expression in the islets of T1D individuals^18^, and demonstrate a preferential upregulation of HLA-B over HLA-A.

### Islet-infiltrating CD8^+^ T-cells from HLA-B40^+^ T1D patients recognize HLA-B40-restricted β-cell peptides

The next question was whether this preferential HLA-B upregulation in insulitis lesions translated into recognition of HLA-B-restricted peptides by islet-infiltrating T-cells. To this end, we first screened 101 T-cell receptors (TCRs), sequenced from islet-infiltrating CD8^+^ T-cells of 4 HLA-B40^+^ T1D organ donors^32^ from nPOD (Supplementary Table 2) and re-expressed into ZsGreen-NFAT fluorescent reporter 5KC T-cells^33^, against 29 HLA-B40-restricted peptides derived from granule proteins and detected in IFN-α-treated β-cells (Supplementary Table 3). We identified a low-affinity TCR 173.D12 that recognized weakly but reproducibly a PPI_44-52_ peptide (Fig. 6A). We reasoned that the preferential HLA-B upregulation should translate into a preferential boosting of HLA-B-restricted T-cell activation upon exposure to IFN-treated β-cells. We therefore compared the ZsGreen-reported activation of HLA-B40-restricted PPI_44-52_ 5KC T-cells with that of HLA-A2-restricted PPI_15-24_ counterparts upon exposure to ECN90 β-cells (Fig. 6B). HLA-A2-restricted T-cells displayed higher activation than HLA-B40-restricted ones when exposed to untreated β-cells, which reflects their higher affinity (Fig. 6A) and the higher abundance of PPI_15-24_ presentation (Fig. 6C). However, HLA-A2-restricted T-cell activation was only marginally increased upon exposure to β-cells treated with IFN-α or IFN-γ, even when adding exogenous peptide to normalize antigen exposure. Conversely, HLA-B40-restricted T-cells were poorly, if at all, stimulated by untreated β-cells, although some potentiation was observed with IFNs. When normalizing antigen presentation by PPI_44-52_ peptide pulsing, the IFN-mediated enhancement was readily visualized. Both T-cells exposed to *INS* knock-out (KO) ECN90 β-cells did not show any reactivity, irrespective of prior β-cell treatment, confirming their antigen specificity. Also in this case, peptide pulsing enhanced HLA-B40-restricted T-cell activation to a larger extent.

**Figure 6.**
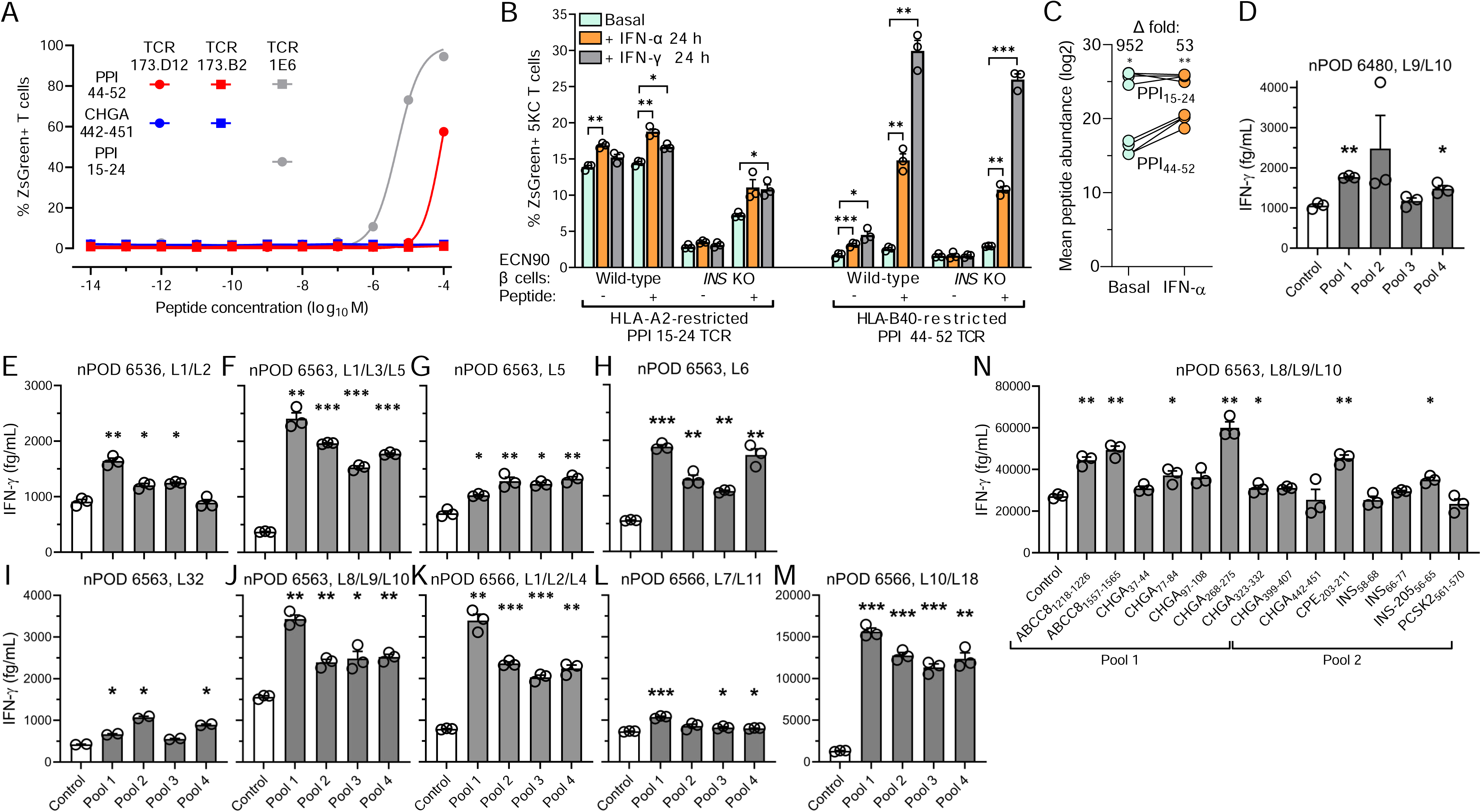
Recognition of HLA-B40-restricted peptides in the islets of T1D donors. **A.** Dose-response peptide recall of 173.D12, 1E6 and negative control 173.B2 TCR-transduced ZsGreen-NFAT reporter 5KC T-cells co-cultured for 18 h K562 antigen-presenting cells transduced with HLA-B40 (for 173.D12 and 173.B2) or HLA-A2 (for 1E6) and pulsed with the indicated peptides. A representative experiment out of 2 performed is shown. **B.** Activation of ZsGreen-NFAT reporter 5KC T-cells transduced with a 1E6 TCR recognizing HLA-A2-restricted PPI_15-24_ or a 173.D12 TCR recognizing HLA-B40-restricted PPI_44-52_. Following the indicated cytokine pretreatment, ECN90 β-cells (wild-type or *INS* KO) left unpulsed or pulsed with the cognate peptide were put in contact with TCR-transduced 5KC T-cells for 6 h. Data represent mean±SEM of triplicate measurements from a representative experiment performed in triplicate. \**p*<0.05, \*\**p*<0.01 and \*\*\**p*<0.001 by Student’s t test. **C.** Average total abundance and fold difference of HLA-A2-restricted PPI_15-24_ and HLA-B40-restricted PPI_44-52_ peptides presented under basal and IFN-α-treated conditions (n=4/each; PPI_15-24_ but not PPI_44-52_ was detected in 4 additional replicates). \**p*<0.05 and \*\**p*<0.01 by paired Student’s t test. **D-N.** IFN-γ secretion by polyclonal CD8^+^ T-cell lines expanded from islet infiltrates of HLA-B40^+^ nPOD T1D donors (listed in Supplementary Table 2) and exposed to HLA-B40-transduced K562 antigen-presenting cells pulsed with HLA-B40-restricted peptide pools (D-M; listed in Supplementary Table 3) or with individual peptides (N). Data represent mean±SEM of triplicate measurements from a representative experiment performed in duplicate. \**p*<0.05, \*\**p*<0.01 and \*\*\**p*<0.001 by paired Student’s t test.

Last, we looked for further evidence of HLA-B40-restricted responses in CD8^+^ T-cell lines raised from islet infiltrates of 4 HLA-B40^+^ nPOD T1D donors^34^ (Supplementary Table 2). The previous 29 HLA-B40-restricted peptides were first screened by combining them into 4 pools (Supplementary Table 3), using peptides binding to irrelevant HLA-I alleles not expressed by donors as negative controls (Supplementary Table 4). All 4 donors yielded some T-cell lines positive to most peptide pools. Donor 6480 displayed positive IFN-γ secretion responses in pooled lines (L)9/L10 (Fig. 6D), while 3 other pooled T-cell lines were negative (not shown). Donor 6536 tested positive for pooled L1/L2 (Fig. 6E); two other pooled lines and individual L6, L9 and L32 were instead negative (not shown). Donor 6563 yielded robust IFN-γ responses in pooled lines L1/L3/L5 (Fig. 6F). L5 was also tested individually, yielding lower yet significant responses (Fig. 6G). L6 from the same donor also responded robustly (Fig. 6H), while L32 responded weakly (Fig. 6I). Pooled L8/L9/L10 gave robust IFN-γ responses to all 4 peptide pools, notably to pool 1 (Fig. 6J). Donor 6566 displayed robust responses to all 4 peptide pools in pooled L1/L2/L4 (Fig. 6K) and L10/L18 (Fig. 6M), minor reactivities for pooled L7/L11 (Fig. 6L) and no reactivity for pooled L2/L3/L4, L12/L14/L17 and individual L6 (not shown). Overall, reactivities were found in 10/10 positive T-cell lines for peptide pool 1, 8/10 for pool 2, 8/10 for pool 3, and 9/10 for pool 4. The reactivity of positive CD8^+^ T-cell lines L8/L9/L10 from donor 6563 was subsequently deconvoluted by testing for each individual peptides of the 4 positive pools (Fig. 6N). While deconvolution of pools 3-4 did not return any hit (not shown), significant IFN-γ secretion was detected for 5/7 peptides from pool 1 (ABCC8_1218-1226_, ABCC8_1557-1565_, CHGA_77-84_, CHGA_268-275_, CHGA_323-332_) and 2/7 peptides from pool 2 (CPE_203-211_, INS-205_56-65_).

Collectively, these data indicate that multiple HLA-B*40-restricted peptides are targeted by islet-infiltrating T-cells from HLA-B40^+^ T1D donors.

## Discussion

Our study provides an in-depth view of the repertoire of HLA-I-bound peptides presented by β-cells exposed or not to the early T1D signature cytokine IFN-α. Expectedly, IFN-α exposure increased HLA-I surface expression^4,19^, which resulted into a higher number and abundance of presented peptides, as we previously reported for IFN-γ^20^. Thus, the inflammatory milieu of insulitis likely enhances the antigenic visibility of β-cells, both qualitatively and quantitatively. These peptides originated from a wider array of source proteins, possibly reflecting IFN-α-induced ER stress^4^ and increased numbers of mis-folded proteins undergoing proteasomal degradation^24^. Within this more diversified peptide display, the fraction derived from secretory granule proteins decreased. However, this decreased fraction reflected a dilutional effect, because both the number and abundance of granule-derived peptides concomitantly increased, underlining the major contribution of secretory granules to the pHLA-I display of β-cells^20,35^ under basal and, to a larger extent, IFN-α-treated conditions. In line with previous findings^36^, de-differentiation, which could have resulted in increased degradation of granule proteins and presentation of derived peptides, was not at play, as IFN-α rather increased the expression of β-cell identity genes such as *INS* and *CHGA*. Accordingly, proINS synthesis was also unaffected by IFN-α, while it was decreased by IFN-γ. β-cell de-differentiation might thus be a late event in T1D pathogenesis, induced by inflammatory cytokines that intervene later than IFN-α^37^, and may represent a defense mechanism to limit autoimmune vulnerability^38^. At earlier disease stages, the HLA-E upregulation induced by IFN-α^19^ may provide another line of defense. HLA-E inhibits NK-cell-mediated cytotoxicity^39^ and reportedly presents a limited set of peptides to regulatory CD8^+^ T-cells^40,41^. Accordingly, the peptides predicted to bind exclusively to HLA-E were presented only by IFN-α-treated β-cells and will provide a useful resource for follow-up studies.

The immunopeptidome obtained confirmed our previous findings^20^ that some known β-cell antigens are underrepresented (ZnT8, GAD65, IAPP; with IAPP possibly reflecting a model bias, as ECN90 express low IAPP levels^26^) or even absent (IGRP, INS DRiP), despite a much deeper peptide sequencing depth. Indeed, using higher cell numbers and more restrictive bioinformatics pipelines, we retrieved 714 conventional 8-12mer peptides as compared to the previous 78^20^. This does not exclude that these missing antigens may become T-cell targets upon peptide presentation by antigen-presenting cells phagocytosing β-cell material. The INS DRiP case is noteworthy, as it has been confirmed as a relevant T-cell antigen targeted by islet-infiltrating CD8^+^ T-cells^32^. The short-lived and unstable nature of INS DRiP and the fact that its presentation is enhanced by IFN-γ but not IFN-α^42^ may explain this discrepancy, emphasizing the need for complementary antigen identification strategies^43^. The proteasome-independent generation of some peptides identified may reflect their processing in other cell compartments. One notable example is PPI_15-24_, which is reportedly processed by signal peptidases during the import of nascent proINS into the ER^44^. We also identified novel antigens, namely CHGB, SCG2 and SCG3, which share several features with the previously reported SCG5, UCN3 and PCSK2^20,35^. They are all soluble granule proteins undergoing intermediate processing by proconvertases and furins to yield their bioactive products, which are released with secretory granules. This processing, along with the modulation of their biosynthesis according to metabolic demand and inflammatory context, may easily divert them toward the HLA-I pathway under β-cell stress conditions^24^.

The comparison of the immunopeptidome of ECN90 β-cells and primary human islets was also informative. Several peptides restricted for the shared HLA-A2/A3 alleles identified in ECN90 β-cells were also found in islets, lending support to this model. Intriguingly, several GCG-derived peptides were also retrieved from islets (n=43 vs. 33 for INS), with some of them mapping to the signal sequence, as for INS (Supplementary Fig. 7). Moreover, α-cells from T1D patients displayed an HLA-I hyper-expression equivalent to that of β-cells compared to T1D controls. These findings suggest that the autoimmune resistance of α-cells cannot be ascribed to a lesser antigen presentation leading to immune ignorance. Other α-cell-intrinsic defense mechanisms and/or a higher proneness of the T-cell repertoire to recognize INS rather than GCG^32,45^ may be at play. Interestingly, one of the GCG-derived peptides mapping to the signal sequence (GCG_2-10_) displays a predicted HLA-E restriction, and α-cell reportedly express higher HLA-E levels than β-cells^45^.

Our thorough investigation of HLA-eluted peptide neo-sequences led to the robust identification of 1 HLA-B40/B49-restricted INS-205 mRNA splice peptide; 9 *cis*-spliced candidates and 12 sequences bearing PTMs. Among these PTMs, citrullination was not found, despite being a relevant PTM in T1D^46–48^. However, citrullinated arginine has the same mass increase as deamidated glutamine or asparagine, which makes their MS distinction challenging^49^. Citrullination may thus be mis-identified as deamidation in peptides containing both an arginine and a glutamine or asparagine. To exclude this possibility, we focused on the 5/13 spectra of arginine-containing deamidated sequences that did not match those of synthetic deamidated peptides and synthesized them in citrullinated form. Also in this case, no spectral matching was observed, thus ruling out the presence of citrullinated peptide. It remains possible that citrullinated epitopes may be generated outside β-cells. The most represented PTM (8/12 peptides) was S-glutathionylation, which involves the formation of a disulfide bond between a cysteine residue and a glutathione tripeptide. Glutathione is a key circulating anti-oxidant, whose levels are reduced in T1D patients due to increased utilization^50,51^. It participates to INS degradation by reducing the disulfide bonds between the A and B chain^52^. The observation that more HLA-I-bound glutathionylated peptides are found in IFN-α-treated conditions may reflect the altered redox balance of inflamed β-cells^53^, a therapeutic target that yielded promising results in recent trials^54^. Moreover, epitope glutathionylation can alter T-cell recognition^55^, implying that oxidative stress may also contribute to the antigenic visibility of β-cells. Cysteines are exquisitely sensitive to oxidative PTMs, with cysteinylation (i.e. a disulfide-linked cysteine addition) also found in our dataset and previously reported in HLA-I-restricted T-cell epitopes from other fields^56,57^. Most PTM peptides (11/12), *cis*-spliced peptides (6/9) and the INS-205 mRNA-spliced peptide were found exclusively after IFN-α treatment, underlining the role of IFN-α in shaping this neo-antigen landscape. This neo-antigen generation may further increase the autoimmune vulnerability of β-cells.

Besides HLA-I hyper-expression and neo-antigen generation, preferential HLA-B upregulation by IFN-α may further contribute to the increased antigenic visibility of β-cells. This effect is not unique to β-cells or IFN-α, as it has been reported for IFN-γ in cancer cells^21^. HLA-B have also been found enriched in extracellular vesicles, along with its peptide ligands^58^. Also in our case, preferential HLA-B upregulation translated into an increased peptide presentation drastically skewed toward HLA-B ligands. Although remaining unnoticed, a preferential upregulation of HLA-B and, to a lesser degree, HLA-C, was also observed in RNA-seq studies of induced pluripotent stem cell-derived islet-like cells exposed to IFN-α^23^; of sorted β-cells from T1D vs. non-diabetic donors^53,59^, despite a long disease duration; and in proteomics studies of islets from non-diabetic donors exposed to IFN-γ/IL-1β^60^. Mechanistically, this may reflect the structure of the *HLA-B* gene locus, which harbors two IFN response elements, while HLA-A and -C loci have only one^21,61^. Moreover, HLA-B ligands originated from a very restricted set of source proteins, with CHGA and INS accounting together for 39% and 11% of the total number of peptides displayed under basal and IFN-α-treated conditions, respectively, as compared to 7% and 4% for HLA-A. This restricted set may also reflect the fact that the ECN90 β-cell model limited our analysis to HLA-B40/B49-binding peptides, whose near-exclusive preference for glutamic acid at position 2 significantly reduces the number and amount of binders. Despite this constraint, the activation of HLA-B40-restricted PPI_44-52_-reactive T-cells was boosted when β-cells were exposed to IFNs, while the activation of HLA-A2-restricted PPI_15-24_-reactive T-cells was only marginally increased. Moreover, islet-infiltrating CD8^+^ T-cells from T1D patients recognized HLA-B40-restricted granule peptides, notably derived from the known β-cell antigen CHGA and the newly described ATP-binding cassette subfamily C member 8 (ABCC8) and carboxypeptidase E (CPE). Given the different peptide-binding preferences of each HLA-I type, this implies that the inflammatory microenvironment of insulitis may skew the autoimmune response toward a distinct set of HLA-B-restricted peptides and recruit additional CD8^+^ T-cell clonotypes. This finding is also relevant in view of the paucity of HLA-B-restricted β-cell epitopes described^62^; and of the notion that HLA-B*39:06 is the strongest T1D-predisposing HLA-I allele (relative risk 5.6)^63^. Moreover, it has recently been reported that HLA-B matching, but neither HLA-A nor HLA-DR matching, improves islet allograft survival^64^. This novel paradigm of preferential HLA-B upregulation may also apply to other autoimmune diseases featuring IFN signatures^65^.

This study carries limitations. First, our work focused on predicted HLA-I ligands, and the datasets used to train prediction algorithms are mostly derived from viral and tumor immunology studies. While in these studies HLA-I binding is a good predictor of T-cell immunogenicity, it likely underestimates the number of potential epitopes in the case of autoimmunity^43^, possibly reflecting a bias imposed by thymic T-cell deletion. Predicted non-binders may hide additional immunogenic peptides, as recently demonstrated for cancer neo-epitopes^66,67^. Second, we analyzed T-cell recognition in only a subset of HLA-B40-restricted peptides. The large repertoire of candidate epitopes identified will require higher-throughput technologies based on DNA-barcoded HLA-I multimers^68^ for a comprehensive validation.

In conclusion, our study shows that IFN-α shapes the immunopeptidome of β-cells, promoting neo-antigen formation and drastically increasing the presentation of HLA-B-restricted peptides. Islet-infiltrating CD8^+^ T-cells recognize a subset of these peptides derived from granule proteins. This comprehensive catalog of HLA-I-eluted peptides sheds light on a neglected, distinct set of peptides presented by HLA-B molecules and invites further studies to identify diabetogenic HLA-B-restricted CD8^+^ T-cells and T-cell biomarkers.

## Methods

### β-cell culture and treatments

The ECN90 β-cell line^33^ was maintained in DMEM/F12 Advanced medium (ThermoFisher) supplemented with 2% bovine serum albumin (BSA; fraction V, fatty-acid-free; Roche), 50 µM β-mercaptoethanol (Sigma), 10 mM nicotinamide (Merck), 1.7 ng/mL sodium selenite (Sigma), 100 U/mL penicillin-streptomycin (Gibco), 1% GlutaMAX (Gibco). Cells were seeded at 8-8.5×10^6^ cells in 75 cm² or 16.5×10^6^ cells in 150 cm² culture flasks (TPP) coated with 0.25% fibronectin from human plasma (Sigma) and 1% extracellular matrix from Engelbreth-Holm-Swarm murine sarcoma (Sigma) and cultured at 37°C in 5% CO_2_ for 18-24 h. ECN90 cells were treated in DMEM/F12 medium (ThermoFisher), supplemented as above or without BSA with the following agents and final concentrations: IFN-α (PBL Assay Science, #11100-1; 2,000 U/mL), IFN-γ (RnD, #285-IF-100, 500 U/ml), carfilzomib (Selleckchem, #S2853; 50 nM) or ONX-0914 (Selleckchem, #S7172; 100 nM). The proteasome enzymatic activity was measured using Proteasome-Glo (Promega #G1180) on ECN90 β-cells collected by trypsinization and extensively washed with PBS.

Human pancreatic islets were obtained from 2 non-diabetic brain-dead organ donors (79-year-old male, BMI 27.8 kg/m², insulin secretion 25.2 and 51.0 µU/ml at 3.3 and 16.7 mM glucose, respectively; 80-year-old female, BMI 21.6 kg/m², insulin secretion 19.6 and 29.6 µU/ml at 3.3 and 16.7 mM glucose, respectively); protocol approved by the Ethics Committee of the University of Pisa, Italy.

The *INS* KO ECN90 β-cell line was generated by transfection with Lipofectamine CRISPRMAX (Invitrogen, #CMAX00001) with Alt-R S.p. Cas9 Nuclease V3 (IDT, #1081058) and *INS*-targeting gRNA (IDT). *INS* KO was validated by RT-PCR and by INS ELISA (Mercodia, #10-1113-01) according to the manufacturer’s protocol.

### HLA immunoprecipitation and peptide elution

Purified anti-HLA-I Ab W6/32 (8-16 mg; produced in-house) was incubated with protein A Sepharose beads for 30 min before being washed with borate buffer (0.05 M boric acid, 0.05 M KCl, 4 mM NaOH, pH 8.0). Bound Abs were cross-linked to the beads with 40 mM dimethyl pimelimidate dihydrochoride (Sigma) in borate buffer pH 8.3 for 30 min. The cross-linking was terminated with ice-cold 0.2 M Tris pH 8.0, and unbound Abs removed by washing with 0.1 M citrate buffer pH 3.0 followed by 50 mM Tris pH 8.0.

Dry-frozen cell pellets (1×10^8^) were lysed in 1 mL lysis buffer (1% IGEPAL-CA 630, 300 mM NaCl, 100 mM Tris pH 8.0, 1X Roche cOmplete Mini Protease Inhibitor Cocktail, EDTA-free) by mixing at 4°C for 30 min. Human primary islets were lysed in lysis buffer in bead beater tubes (Precellys Evolution, Bertin) following 5 cycles at 7,200 rpm for 20 s, separated by 20 s pauses. Lysates were collected, further mixed for 20 min at 4°C, and cleared by centrifugation at 500 g for 10 min followed by 21,000 g for 45 min at 4°C. pHLA complexes were captured by incubating the cleared lysates with W6/32 Ab-cross-linked Sepharose A beads overnight at 4°C. The resin was collected by gravity.

### HPLC fractionation and purification of HLA-I-bound peptides

ECN90 β-cell samples were resuspended in 120 µL loading buffer (0.1% TFA, 1% acetonitrile) and injected in an Ultimate 3000 HPLC System (ThermoFisher). Peptides were separated across a 4.6-by 50-mm ProSwift RP1S column (ThermoFisher) using 1mL/min flow rate over 10 min from 2% to 35% buffer B (0.1% TFA in acetonitrile) in buffer A (0.1 TFA in water). Fifteen fractions were collected, every 30 s, and peptide fractions 1-9 were combined into odd and even aliquots and vacuum-dried prior to LC-MS/MS acquisition.

Resuspended HLA-I-eluted samples were centrifuged through 5 kDa cutoff filters (Merck Millipore #UFC3LCCNB-HMT) and vacuum-dried. They were then resuspended in loading buffer and cleared using Pierce C18 Spin Tips (ThermoFisher, #84850). Final elution was performed in 30% acetonitrile 0.1% TFA. Samples were vacuum-dried prior to LC-MS/MS acquisition.

### LC-MS/MS acquisition

Dried samples of HLA-I-bound peptides from ECN90 β-cells exposed or not to IFN-α and synthetic peptides were resuspended in loading buffer (1% acetonitrile, 0.1% TFA) and analyzed by an Ultimate 3000 RSLCnano system coupled to a Fusion Lumos mass spectrometer (ThermoFisher). Peptides were separated using a PepMap C18 column, 75 µm × 50 cm, 2 µm particle size (ThermoFisher) with a 30 min (synthetic peptides) to 60 min linear acetonitrile in water gradient of 2-25%, at a flow rate of 250 µL/min. Solvent contained 5% DMSO and 0.1% formic acid (v/v). Peptides were ionized using an EasySpray source at 2,000 V and ions were introduced into the mass spectrometer through an on-transfer tube at 305°C. Data-dependent acquisition was performed with one full MS1 spectra recorded from 300 to 1,500 m/z (120,000 resolution, 400,000 AGC target, 60 ms accumulation time), followed by MS2 scans (30,000 resolution, 120 ms accumulation, 300,000 AGC target). Precursor selection was performed using TopSpeed mode at a cycle time of 2 s. High-collision dissociation (HCD) fragmentation was induced at an energy setting of 32 for singly charged peptides and of 28 for peptides with a charge state 2-4.

Alternatively, vacuum-dried samples of HLA-I-bound peptides from ECN90 β-cells exposed to (immuno-)proteasome inhibitors or human islets were resuspended in SCP loading buffer (1% acetonitrile, 0.1% formic acid) and analyzed by a nanoElute system coupled to a timsTOF SCP mass spectrometer (Bruker). Peptides were loaded into an Aurora C18 column, 25 cm × 75 µm, 1.7 µm particle size (IonOpticks) with a loading pressure of 800 bars for 9 min and were separated with a linear acetonitrile gradient of 2-25% in 0.1% acetic acid over 60 min and 24-37% for additional 6 min at a flow rate of 150 nL/min at 50°C. Peptides were ionized with the CaptiveSpray source (Bruker) at 1,400 V and 180°C. Data were acquired in DDA PASEF mode with one TIMS-MS survey and 10 PASEF MS2 scans per cycle. Ion accumulation and ramp time in the TIMS analyzer were set to 166 ms/each. The ion mobility range for peptide analysis was set to 1/K0 = 1.7 to 0.7 Vs/cm² and the m/z range was 100-1,700. Two compound regions were defined using m/z and 1/K0 Vs/cm² as follows: 300, 0.7; 800, 1.2; 800, 0.9; 500, 0.7; and 700, 1.4; 700, 1.1; 1000, 1.7; 1500, 1.7. Precursors with charge states 1-3 and a minimum threshold of 500 arbitrary units (AU) were fragmented and re-sequenced until reaching a “target value” of 20,000 AU. Collision energies were 70 eV at 1/K0 = 1.7 Vs/cm^2^; 40 eV at 1/K0 = 1.34 Vs/cm^2^; 40 eV at 1/K0 = 1.1 Vs/cm^2^; 30 eV at 1/K0 = 1.06 Vs/cm^2^; 20 eV at 1/K0 = 0.7 Vs/cm^2^.

### LC-MS/MS data analysis

Analysis of the raw data was performed using PEAKS X or PEAKS X Pro (Bioinformatics Solutions), reporting 5-10 peptides for each spectrum in *de-novo* sequencing, and searching a protein sequence FASTA file containing the reviewed human Uniprot entries (downloaded on 22/01/2019), predicted peptide neo-sequences translated from published RNAseq datasets^69^ and previously reported INS DRiPs^25^. PEAKS PTM search was performed with all 314 built-in modifications. Some specific PTM searches were run by setting the PTM of interest as a variable modification in both the *de-novo* and database search. The false discovery rate (FDR) was calculated with a decoy database search integrated into PEAKS and set to 1% for all samples. A 5% FDR was allowed when looking for mRNA variants reported in human islets (see *Immunopeptidomics bioinformatics analysis*). Label-free quantification of peptide abundance was performed using Progenesis QI (Waters) for chromatographic alignment, normalization, and determination of ion abundances based on the area. Data analysis was performed with Python, Perseus and Excel. Sequence clustering was generated by GibbsCluster (https://services.healthtech.dtu.dk/). All other figures and statistical analyses were done using GraphPad Prism (v9.0). Datasets are available via ProteomeXchange with identifiers PXD045265 and PXD045211.

### RNAseq datasets and analysis

RNAs from 6 individual preparations of primary human islets exposed or not to IFN-α for 18 h^19^ and from HLA Class II^lo^ and HLA Class II^hi^ human medullary thymic epithelial cells (mTECs)^20^ were sequenced on an Illumina HizSeq 2000 at high depth (coverage >150×10^6^ reads, which is sufficient to detect >80% splice variants). Gene expression was quantified using Salmon version 0.13.2^70^ with parameters “--seqBias –gcBias –validateMappings”. GENCODE v31 (GRCh38)^71^ was chosen as the reference genome and has been indexed with the default k-mer values. The estimated number of reads obtained from Salmon were used as input to perform differential expression with DESeq2 1.24.0^70^. For each gene included in DESeq2’s model, a log2 fold change (FC) was computed and a Wald test statistics was assessed with unadjusted and adjusted *p* values. Transcripts were considered differentially expressed when presenting a FC >1.50 and an adjusted *p* value <0.05. Only transcripts presenting >0.5 transcripts per million (TPM) in at least 20% of samples were selected for further analysis. Datasets have been deposited under GEO: GSE148058.

For the RNAseq pipeline, 30,947 mRNAs (TPM >0.5) were filtered based on:

a) A median TPM>5 in islets, either under basal or inflammatory conditions, a cut-off selected based on the median TPM of known islet antigens (islet expression filter; n=12,594).
b) A median TPM<0.1 in mTECs (either HLA Class II^lo^ or HLA Class II^hi^), or a median TPM fold-decrease >100 vs. islet (mTEC expression filter).
c) A median TPM fold-increase >10 in islets compared to 12 control tissues (adipose tissue, breast, colon, heart, kidney, liver, lung, lymph node, ovary, prostate, skeletal muscle, white blood cells), using the Illumina BodyMap 2.0 dataset (islet enrichment filter). Tissues of neuroendocrine origin (brain, testis, adrenal gland and thyroid) were excluded for this filtering.
d) We subsequently focused our analysis on mRNA isoforms, as described^20^. The predicted translation products were aligned using the R package Biostrings v2.52.0, with BLOSUM100 as the score matrix, and aa neo-sequences were defined by comparing the predicted aa sequence of each mRNA isoform with that of the reference (canonical) mRNA, taking as reference the longest and/or most prevalent mRNA isoform in islets (neo-sequence generation filter).

### Immunopeptidomics bioinformatics analysis

An in-house analysis pipeline was designed (Python 3.7) and applied to the PEAKS PTM output file. After discarding long (>14-aa) sequences, the remaining ones were sorted based on their source proteins/genes and according to their expression in the pancreas, based on downloadable databases and datasets. Briefly, peptides whose source genes were found in the pancreas at the RNA level, or at the protein level with a high/medium degree of evidence according to the Human Protein Atlas (V18) or the Human Protein Reference Database (release 9) were retained. Previously published single-cell RNAseq datasets from human pancreatic islets^72^ were re-processed to generate a list of genes that are enriched in endocrine cells. Peptides identified in MS were further retained only if their source gene was found in this list. Together, these steps defined the “β-cell-enriched expression” filter. In parallel, the spectra found in the PEAKS PTM search (matched to the database) were excluded from the PEAKS *de-novo* search output, and the remaining spectra were searched for *cis*-spliced peptide sequences. The 10 reported peptides per spectrum were investigated for putative forward (i.e. ligation of two fragments in the order in which they occur in the parent protein) or backward (i.e. ligation of two fragments in the reverse order compared to the parent protein sequence) *cis*-spliced sequences of proteins included in the reference database. Sequences were then fed into the MARS v1.0 algorithm^27^, trained with the genome-templated sequences. The MARS output and the putative *cis*-spliced sequences were compared, and sequences found in both were retained and filtered based on their β-cell-enriched expression as above. NetMHCpan4.1a (https://services.healthtech.dtu.dk/) was installed locally and used to predict HLA-binders to the alleles expressed by ECN90 β-cells, i.e. HLA-A*02:01, -A*03:01, -B*40:01, -B*49:01, -C*03:04, -C*07:01 and -E*01:01 (cutoff score of <2)^28^. For PTM peptides, the prediction was performed on native sequences. HLA-I binding predictions were not computed for *cis*-spliced sequences, as netMHCpan ranking is a selection parameter already embedded in the MARS algorithm. A single HLA prediction was attributed to a peptide if the score difference with the second-best allele was ≥3-fold. Otherwise, the two best HLA prediction were assigned.

### Real-time quantitative PCR (RT-qPCR)

Total RNA was isolated from ECN90 β-cells using RNeasy mini kit (Qiagen), with RNA concentration and purity assessed by Nanodrop (ThermoFisher). cDNA was synthesized from 500 ng total RNA using superscript VILO synthesis kit (ThermoFisher #11754050) prior to gene expression analysis by real-time quantitative PCR (LightCycler, Roche; or QuantStudio 3, ThermoFisher). Quantitect SYBR green PCR mastermix (Qiagen) or Power SYBR green mix (Applied Biotechnologies) along with primers designed with Primer3 software were used (Supplementary Table 5). For HLA-I genes, primers were as reported^21^. Melting curve analysis was performed to evaluate the specificity of each amplicon prior to analysis. To normalize gene expression across samples, *GAPDH*, *ACTB* and *PPIA* were used as housekeeping genes. The ΔCt comparative method for relative quantification was used.

### Protein synthesis analysis by puromycin incorporation

IFN-α-or IFN-γ-treated ECN90 β-cells were pretreated or not with cycloheximide (50 μg/mL; Sigma) for 1 h before pulsing during the last 10 min with puromycin (10 μg/mL; ThermoFisher). After 3 washes with ice-cold PBS, cells were harvested, paraformaldehyde-fixed and stained with AF488-coupled anti-puromycin Ab (RRID:AB_2736875) and APC-coupled anti-HLA-A/B/C/E Ab (RRID:AB_314879) in PBS/BSA with 1% saponin (ThermoFisher #C10424) prior to acquisition on a BD LSRFortessa.

Neo-synthetized puromycin-labeled proINS was analyzed in ECN90 β-cells lysed in RIPA buffer with protease and phosphatase inhibitors. Proteins (600 µg) were incubated with proINS Ab RRID:AB_10949314 overnight at 4°C, followed by addition of protein A/G agarose beads (ThermoFisher) at room temperature for 3 h. Immunoprecipitated complexes were washed 4 times and eluted using Laemmli buffer (Bio-Rad). Low-molecular-weight protein Western blot was performed as described^73^. PVDF membranes were incubated with anti-puromycin Ab overnight at 4°C and secondary Ab RRID:AB_330924 at room temperature for 2 h and revealed by SuperSignal West Dura and iBright imaging system (ThermoFisher). After stripping, membranes were incubated with proINS Ab overnight at 4°C and processed as above.

### Analysis of HLA-A, HLA-B and HLA-C expression

For Ab validation, HLA-I^−^ K562 cells transduced with different HLA-I alleles^32,33^ were single-stained with the following Abs: HLA-A clone ARC0588 (RRID: AB_2849011) with secondary Ab RRID:AB_2536097; HLA-B clone JOAN-1 (RRID:AB_1076708) with secondary Ab RRID:AB_2536161; HLA-C Ab DT-9 (RRID:AB_2739715); HLA-A/B/C/E Ab W6/32 (RRID:AB_314873), followed by acquisition on a Beckman CytoFLEX flow cytometer. Surface HLA-I expression was analyzed on ECN90 β-cells single-stained for 30 min with Live/Dead Violet (ThermoFisher) and Abs to: HLA-A clone ARC0588 (RRID: AB_2849011) with secondary Ab RRID:AB_2832926; HLA-B clone JOAN-1 (RRID:AB_1076708), HLA-C Ab DT-9 (RRID:AB_2650941), HLA-A/B/C/E Ab W6/32 (RRID: AB_314871) with secondary Ab RRID:AB_2921066 before acquisition on an LSRFortessa and analysis with FlowJo v10.8.

For Western blotting, cell pellets were lysed in RIPA buffer with protease inhibitor cocktail (Sigma) and total protein was quantified with Pierce BCA protein assay kit (ThermoFisher). Protein samples were separated by Bolt 4–12% Bis-Tris polyacrylamide gels (Invitrogen) under denaturing conditions. Detection was performed using the Abs ARC0588 (HLA-A, RRID:AB_2849011; 1/1,000) and HC10 (HLA-B, RRID:AB_2728622; 1/1,000) overnight at 4°C, followed by horseradish peroxidase-conjugated secondary Abs RRID:AB_2687483 and RRID:AB_330924 (1/10,000), respectively. α-tubulin Ab RRID:AB_1210457 (1/1,000, 1 h at room temperature) was used with RRID:AB_330924 for loading control. Bands were quantified with ImageJ.

For multiplex tissue immunofluorescence, formalin-fixed paraffin-embedded pancreas tissue sections were obtained from EADB (https://pancreatlas.org/; with ethical permission from the West of Scotland Research Ethics Committee, 15/WS/0258) or nPOD (Supplementary Table 1). Sections were baked at 60°C for 1 h, dewaxed in Histoclear, rehydrated in degrading ethanol concentrations (100%, 95%, 70%) and fixed in 10% neutral-buffered formalin. Heat-induced epitope retrieval (HIER; 10 mM citrate, pH 6) was performed for epitope unmasking by placing sections in a pressure cooker in a microwave oven at full power for 20 min. The sections were then blocked with 5% normal goat serum and incubated with primary Ab, followed by probing with an appropriate OPAL fluorophore-conjugated secondary Ab (Akoya Biosciences, Supplementary Table 6). This was followed by a further HIER to remove the primary Ab before staining with the next primary/secondary Ab combination. The same steps (from blocking to epitope retrieval) were repeated 5 times (for each of the 6 primary/secondary Ab combinations). Sections were counterstained with DAPI and mounted for multispectral fluorescent microscopy using the Vectra Polaris slide scanner (Akoya Biosciences). Quantification was performed using the Indica HALO image analysis platform. The DenseNet classification module was used to identify endocrine and exocrine regions on the whole slide scan. The resolution was set at 2 µm/pixel and the minimum object size at 750 µm^2^. The identified endocrine regions (islets) were manually sorted into ICIs and IDIs. The HighPlex module was used to segment and phenotype islet cells; β-cells were defined as INS^+^GCG^−^ and α-cells as INS^−^GCG^+^. Cell object data were exported in Excel format and analyzed in GraphPad Prism 10.

### Antigen recall on carrier T-cells transduced with TCRs from islet-infiltrating CD8^+^ T-cells

HLA-B40-restricted peptides (Supplementary Table 3) were screened on TCRs obtained from donors listed in Supplementary Table 2. Fluorescent reporter TCR transductants were generated as described^32,33^. Briefly, 5KC T-hybridoma cells carrying transgenes for an NFAT-driven ZsGreen-1reporter and human CD8 were transduced with retroviral vectors encoding TCRs recognizing an HLA-B40-restricted PPI_44-52_ epitope (TCR 173.D12) or an HLA-A2-restricted PPI_15-24_ epitope (TCR 1E6). These TCRs were re-expressed as chimeric TCRα/β pairs linked by a porcine teschovirus-1 2A (P2A) peptide (synthesized by Twist Bioscience). ECN90 β-cells, either wild-type or *INS* KO and pulsed or not with TCR cognate peptides (100 µM, Synpeptide), were exposed for 24 h to IFN-α (PBL, 2,000 U/ml), IFN-γ (RnD, 500 U/ml) or no cytokine prior to addition of TCR transductants (T:β-cell ratio 1:2) for 6 h.

### Antigen recall on islet-infiltrating CD8^+^ T-cells

Isolated islets or live pancreas slices (150-µm thick) were provided by nPOD (donors listed in Supplementary Table 2). Islets were isolated by tissue digestion with 1 mg/mL collagenase-P (Sigma) in PBS at 37°C under shaking with a magnetic stir bar, with tissue dispersal visually monitored. After washing, islets were enriched by hand-picking, and single-islet-derived T-cell lines generated as described^34^. The lymphocytic outgrowth from individual islets was collected, re-plated, allowed to expand, re-stimulated as needed and cryopreserved in early passage (P1-3). T-cell lines containing >20% CD8^+^ T-cells were selected for analysis.

HLA-B40-transduced K562 cells were irradiated (5,000 rads), plated at 3,000 cells/well and pulsed with pools of HLA-B40-restricted peptides (4 pools of 7-8 peptides/each; Supplementary Table 3) or a negative control pool of peptides binding to irrelevant HLA-I alleles (Supplementary Table 4)^44,62^. They were then co-cultured for 48-72 h with individual islet-derived T-cell lines (or pooled lines when needed to reach required T-cell numbers) at 75,000 cells/well in triplicate wells of round-bottom 96-well plates. T-cells alone were cultured with and without plate-bound anti-CD3/CD28 as positive control for functional, viable T-cells. Supernatants were collected and IFN-γ secretion measured by cytometric bead array (BD #561515; 271 fg/ml lower detection limit). The reactivity of pooled T-cell lines L8/L9/L10 from donor 6563 was deconvoluted for individual peptides as above.

### Statistical analysis

Statistical details of experiments can be found in the legends of each figure. A two-tailed p<0.05 cut-off was used to define statistical significance.

## Acknowledgments

We gratefully acknowledge L. Bailly (myBrain Technologies, Paris) for his help in developing the bioinformatics analysis pipeline; S. Salignac (Cochin Institute, Paris) for technical assistance; M. Peakman (King’s College, London) for providing the 1E6 TCR sequence; and the Cochin Institute CYBIO Flow Cytometry Core Facility for assistance with cell analysis. This work was supported by The Leona M. and Harry B. Helmsley Charitable Trust (1901-03689), Agence Nationale de la Recherche (ANR-19-CE15-0014-01) and Fondation pour la Recherche Medicale (EQU20193007831), to R.M.; by the European Foundation for the Study of Diabetes (EFSD/JDRF/Lilly European Programme in Type 1 Diabetes Research 2019), to N.T.; JDRF Postdoctoral Fellowship 3-PDF-2020-942-A-N, to Z.Z.; EFSD/Lilly Young Investigator Research Award Programme 2023, to F.S.; National Institutes of Health (NIH) U01 DK104218-04 and UC4 DK116284, to S.C.K.; NIH R01 DK099317, P30 DK116073, to M.N.; and Steve Morgan Foundation Grand Challenge Senior Research Fellowship (22/0006504), to S.J.R. D.L.E. acknowledges the support of JDRF grants 3-SRA-2022-1201-S-B(1) and 3-SRA-2022-1201-S-B (2), Welbio-FNRS (Fonds National de la Recherche Scientifique; WELBIO-CR-2019C-04), the NIH Human Islet Research Network Consortium on Beta Cell Death & Survival (HIRN-CBDS; U01 DK127786) and NIH/NIDDK grants RO1DK126444 and RO1DK133881-01. R.M., D.L.E., R.S. and S.J.R. received funding from the Innovative Medicines Initiative 2 Joint Undertaking under grant agreements 115797 and 945268 (INNODIA and INNODIA HARVEST), which receive support from the EU Horizon 2020 program, JDRF, and The Leona M. & Harry B. Helmsley Charitable Trust. This research was performed with the support of the Network for Pancreatic Organ Donors with Diabetes (nPOD; RRID:SCR_014641), a collaborative T1D research project sponsored by JDRF (nPOD: 5-SRA-2018-557-Q-R) and The Leona M. and Harry B. Helmsley Charitable Trust (Grant #2018PG-T1D053). Organ Procurement Organizations (OPO) partnering with nPOD to provide research resources are listed at www.jdrfnpod.org/for-partners/npod-partners. The content and views expressed are the responsibility of the authors and do not necessarily reflect an official view of nPOD.

## Author contributions

A.C. performed immunopeptidomics experiments, analyzed data, participated in the conceptualization and supervision of the study, wrote the first draft and edited subsequent drafts. Z.Z. performed HLA-I expression and T-cell transductant experiments and analyzed data. J.P.-H. and O.B.G. performed HLA-I expression experiments and analyzed data. F.S. performed T-cell experiments, analyzed data and participated in the conceptualization of the study. C.L. and S.J.R. conceived and performed pancreas tissue immunofluorescence experiments and analyzed data. A.M. and S.C.K. generated T-cell lines, performed T-cell experiments and analyzed data. M.O. characterized β-cell changes upon IFN treatment and analyzed data. H.L. performed the analysis of *cis*-spliced peptides. R.P. and A.N. participated in immunopeptidomics experiments and data analysis. B.B. helped with the generation and testing of T-cell transductants. M.L.C. and D.L.E. generated the dataset of mRNA splice variants. D.L.E., M.G. and S.R.H. participated in the conceptualization of the study. A.A., L.L. and M.N. participated in the generation of T-cell transductants and validation of HLA-I Abs. F.K. and R.S. contributed wild-type and *INS* KO ECN90 β cells and edited the manuscript. L.M. and P.M. contributed primary human islets. S.Y. contributed to study supervision, funding acquisition and manuscript editing. N.T. supervised the study, provided resources and acquired funding. R.M. conceived and supervised the study, provided resources, acquired funding, and wrote the final version of the manuscript.

