## Supplementary Information for "Interferon-α promotes neo-antigen formation and preferential HLA-B-restricted antigen presentation in pancreatic β-cells"

**Alexia Carré et al.**

**Supplementary Information**

**Supplementary Data 1. HLA-I-eluted peptides derived from mRNA splice variants identified in ECN90 β-cells.** The first column displays the peptide sequence with, in parenthesis, the mass shift of PTMs found on the aa. PTMs are detailed in the next column and were ultimately defined as likely artifactual. Subsequent columns indicate the condition and the number of replicates in which the peptides were identified; aa length; UniProt accession number of the canonical source protein; gene name and description; whether the source is a granule protein (Y, yes; N, no); predicted HLA-I restriction(s) and corresponding ranking based on NetMHC4.1a (NA, not assigned). The last column indicates whether the peptide sequence was validated by spectral matching (ND, not determined).

**Supplementary Data 2. Conventional HLA-I-eluted peptides identified in ECN90 β-cells.** The first column displays the peptide sequence. Subsequent columns indicate the condition and the number of replicates in which the peptides were identified; aa length; UniProt accession number; aa position within the source protein (referring to the first source protein listed); gene name and description; whether the source is a granule protein (Y, yes; N, no); predicted HLA-I restriction(s) and corresponding ranking based on NetMHC4.1a (NA, not assigned).

**Supplementary Data 3. Post-translationally modified HLA-I-eluted peptides identified in ECN90 β-cells.** The first column displays the peptide sequence with, in parenthesis, the mass shift of PTMs found on the aa, with PTMs detailed in the next column. Subsequent columns indicate the condition and the number of replicates in which the peptides were identified; aa length; UniProt accession number; aa position within the source protein (referring to the first source protein listed); gene name, synonyms and description; whether the source is a granule protein; predicted HLA-I restriction(s) and corresponding ranking of unmodified sequences based on NetMHC4.1a; and the final PTM origin assigned as likely artifactual (i.e. found in both the modified and unmodified synthetic peptide) or likely biological (i.e. found only in the modified synthetic peptide). Y, yes; N, no; NA, not assigned; ND, not determined.

**Supplementary Data 4. *Cis*-spliced HLA-I-eluted peptides identified in ECN90 β-cells.** The first three columns indicate the source protein (UniProt accession number and gene name) and whether its expression is β-cell-enriched. Subsequent columns indicate predicted HLA-I restriction(s) and corresponding ranking based on NetMHCpan4.1a; the condition in which the peptides were identified; the full peptide sequence; its N-terminal and C-terminal splicing fragments; the forward or reverse splicing direction; and the inter-fragment distance. The last two columns indicate whether the peptide was ultimately validated by spectral matching; and the Ensembl accession number when mRNA splice isoforms were also assigned by the MARS algorithm as an alternative aa sequence match.

**Supplementary Data 5. HLA-I-eluted peptides identified in primary human islets.** The first column shows the peptide sequence with, in parenthesis, the mass shift of PTMs found on the aa. PTMs are detailed in the next column. Subsequent columns indicate the sample in which the peptides were identified (islet preparation 1 and/or 2); aa length; UniProt accession number of the canonical source protein(s); peptide position in the protein (referring to the first source protein listed); gene name, gene synonyms and description; whether the source protein is a granule protein (Y, yes; N, no); predicted HLA-I restriction(s) and corresponding ranking based on NetMHC4.1a (NA, not assigned). GCG is appended to the list of β-cell-enriched source proteins for comparison, with peptide mapping displayed in Supplementary Figure 7.

**Supplementary Data 6. Predicted HLA-A2- and HLA-A3-restricted peptides identified in ECN90 β-cells and primary human islets.** The comparison was performed between the conventional peptides eluted from ECN90 β-cells and from the two primary islet samples. For PTMs defined as likely artifactual, the native peptide sequence was retained. From left to right, columns indicated: gene name of the source protein, peptide sequence, the treatment condition of ECN90 β-cells in which the peptide was identified, aa length, peptide position in the source protein (referring to the first source protein listed); gene synonyms and description; whether the source protein is a granule protein or not; HLA-I restriction(s) and corresponding ranking based on NetMHC4.1a, and the presence in the immunopeptidome of islet sample 1 and/or 2.

**Supplementary Data 7. PTMs identified in the immunopeptidome of ECN90 β-cells.** The first two columns indicate the PTM name and mass shift observed in MS. The next three columns list the number of peptides and number of source proteins identified with the indicated PTM, along with their distribution between basal and IFN-α-treated conditions. Specific PTM searches were run for those PTMs meeting our selection criteria (shown in bold), i.e. enrichment in either basal or IFN-α-treated condition (barring tryptophan oxidation to kynurenin) and PTMs unlikely to naturally arise during peptide synthesis or MS acquisition and introducible into synthetic peptides. PTMs detected in very few (<10) peptides were only retained if found in predicted HLA-A2 or HLA-A3 binders. In total, 253 PTM sites on 247 peptides (with 6 peptides carrying two distinct PTMs) were identified. The last column indicates the number of peptides for which the PTM was validated by spectral matching, i.e. a match with the peptide synthesized with the modification, but not in the peptide synthesized in its native form. Given their unlikely artifactual origin, phosphorylated and glutathionylated peptides were not compared to the unmodified sequence. Modified peptides validated as likely biological are highlighted in grey.

**Supplementary Data 8. Peptide clusters in ECN90 β-cells exposed to proteasome inhibitors.** The identity of the peptides identified in each of the clusters (first column) depicted in Supplementary Fig. 3C is provided, ranked according to cluster number and gene name.

**Supplementary Data 9. Putative HLA-E*01:01 ligands in the immunopeptidome of ECN90 β-cells.** To assign HLA-E*01:01 restriction, a ≥3-fold NetMHC4.1a rank score difference with the second-best allele was used. This is a less stringent criterion than the one used for assigning HLA-E*01:01 restriction in Supplementary Tables 1-4, namely a predicted HLA-E*01:01 restriction without any other HLA-I allele with a ranking score <2. The list includes both conventional peptides and peptides carrying PTMs (listed as native sequences, as all PTMs found on these peptides were defined as likely artifactual). The first column displays the peptide sequence. Subsequent columns indicate the condition and the number of replicates in which the peptides were identified; aa length; UniProt accession number; aa position within the source protein (referring to the first source protein listed); gene name, synonyms and description; whether the source is a granule protein (Y, yes; N, no); predicted HLA-I restriction(s) and corresponding ranking based on NetMHC4.1a.

| Donor group | nPOD/EADB case RRID | Sex | Age (yrs) | T1D (yrs) | Positive Auto-Abs | C-peptide (ng/mL) |
|---|---|---|---|---|---|---|
| T1D | nPOD 6228 SAMN15879284 | M | 13 | 0 | GAD/IA-2 ZnT8 | 0.10 |
| T1D | nPOD 6247 SAMN15879303 | M | 24 | 0.6 | mIAA | 0.47 |
| T1D | nPOD 6264 SAMN15879318 | F | 12 | 9.0 | None | <0.05 |
| T1D | nPOD 6380 SAMN15879433 | F | 12 | 0 | None | 0.22 |
| T1D | nPOD 6396 SAMN15879449 | F | 17 | 2.0 | None | 0.06 |
| T1D | EADB E560 | F | 42 | 1.5 | ND | ND |
| ND | nPOD 6160 SAMN15879216 | M | 22 | NA | None | 0.40 |
| ND | nPOD 6227 SAMN15879283 | F | 17 | NA | None | 2.75 |
| ND | nPOD 6278 SAMN15879332 | F | 12 | NA | None | 4.54 |
| ND | nPOD 6462 SAMN15879515 | F | 14 | NA | None | 11.09 |

**Supplementary Table 1. nPOD and EADB cases analyzed for the expression of HLA-I.** Pancreas tissue immunofluorescence images and their analyses are presented in Fig. 5 and Supplementary Fig. 6. ND, not determined; NA, not applicable.

| nPOD case RRID:SAMN# | Sex | Age (yrs) | T1D (yrs) | Positive Auto-Abs | HLA-A | HLA-B | HLA-C | Positive/tested TCRs | Positive/tested T-cell lines |
|---|---|---|---|---|---|---|---|---|---|
| 6342<br>15879396 | F | 14 | 2 | IA-2 | 02:01<br>68:01 | **40:01**<br>**40:01** | 03:04<br>03:04 | 0/29 | NA |
| 6480<br>15879533 | M | 17 | 2.5 | IA-2 | 03:01<br>11:01 | 07:02<br>**40:01** | NA<br>NA | NA | 1/4 (pooled) |
| **6536**<br>**18242780** | F | 20 | 4.0 | GAD | 02:01<br>31:01 | 08:01<br>**40:01** | 03:04<br>07:01 | 1/28 | 1/3 (pooled), 0/3 (individual) |
| 6563<br>30386851 | F | 15 | 0 | IA-2 | 02:01<br>11:01 | **40:01**<br>50:01 | 03:04<br>06:02 | 0/20 | 2/2 (pooled), 3/3 (individual) |
| 6566<br>33284286 | M | 16 | 2.0 | GAD/IA-2<br>ZnT8 | 03:01<br>23:01 | **40:01**<br>50:01 | 03:04<br>06:02 | 0/24 | 3/5 (pooled), 0/1 (individual) |

**Supplementary Table 2. nPOD islet donors analyzed for the identification of HLA-B40-restricted TCRs and $CD8^+$ T-cells.** For the identification of HLA-B40-restricted TCRs, TCRs from islet-infiltrating T cells of $HLA\text{-}B40^+$ nPOD donors were sequenced and re-expressed into ZsGreen fluorescent reporter 5KC T cells and tested with peptides listed in Supplementary Table 3. The nPOD case from whom TCR 173.D12 was selected is indicated in bold. For the identification of HLA-B40-restricted $CD8^+$ T cells, islet-derived T-cell lines from $HLA\text{-}B40^+$ nPOD donors were co-cultured with HLA-B40-transduced K562 antigen-presenting cells pulsed with the peptide pools listed in Supplementary Table 3. Pools of peptides binding to irrelevant HLA-I were used as negative controls (listed in Supplementary Table 4). Peptide recognition was assessed by measurement of IFN-γ secretion, as shown in Fig. 6D-N. All T-cell lines tested responded to plate-bound anti-CD3/CD28 stimulation as positive control. The "Positive/tested T-cell lines" column displays the number of positive T-cell lines (tested in pools to reach required T-cell numbers, or individually when possible) out of those tested. NA, not available.

| Pool number | Peptide sequence | Source protein | AA position | Accession number | PTM | Positive samples | HLA restriction | NetMHCPan rank |
|---|---|---|---|---|---|---|---|---|
| **1** | **YEARFQQKL** | **ABCC8** | **1218-1226** | **Q09428** | **0** | **IFN-α** | **B40:01** | **0.0117** |
| **1** | **LEFDKPEKL** | **ABCC8** | **1557-1565** | **Q09428** | **0** | **IFN-α** | **B40:01/B49:01** | **0.0175/0.0366** |
| 1 | VEVISDTL | CHGA | 37-44 | P10645 | 0 | IFN-α | B40:01/B49:01 | 0.357/0.8383 |
| **1** | **KELQDLAL** | **CHGA** | **77-84** | **P10645** | **0** | **IFN-α** | **B40:01/B49:01** | **0.4181/1.154** |
| 1 | HSGFEDELSEVL | CHGA | 97-108 | P10645 | 0 | Both | B40:01 | 0.6446 |
| 3 | VEEPSSKDVM | CHGA | 121-130 | P10645 | 0 | IFN-α | B40:01/B49:01 | 0.5364/1.5396 |
| **1** | **GESRSEAL** | **CHGA** | **268-275** | **P10645** | **0** | **IFN-α** | **B40:01** | **0.4029** |
| **1** | **GELEQEEERL** | **CHGA** | **323-332** | **P10645** | **0** | **IFN-α** | **B40:01** | **0.1737** |
| 2 | REDSLEAGL | CHGA | 399-407 | P10645 | 0 | IFN-α | B40:01 | 0.0618 |
| 3 | LEAGLPLQV | CHGA | 403-411 | P10645 | 0 | Both | B49:01/B40:01 | 0.0173/0.239 |
| 2 | AELEKVAHQL | CHGA | 442-451 | P10645 | 0 | IFN-α | B40:01 | 0.0394 |
| 3 | FEGRELLVI | CPE | 83-91 | P16870 | 0 | IFN-α | B49:01/B40:01 | 0.2671/0.5285 |
| **2** | **KEGGPNNHL** | **CPE** | **203-211** | **P16870** | **0** | **IFN-α** | **B40:01** | **0.0911** |
| 4 | WEDNKNSLI | CPE | 358-366 | P16870 | 0 | IFN-α | B49:01/B40:01 | 0.262/0.4403 |
| 4 | SESC(+305.07)PVVGM | KIF1A | 923-931 | Q12756 | S-glutathionylation | IFN-α | B40:01/B49:01 | 0.212/0.431 |
| **4** | **GERGFFYTP** | **INS** | **44-52** | **P01308** | **0** | **Both** | **B49:01/B40:01** | **0.4123/1.2003** |
| 4 | REAEDLQVGQV | INS | 56-66 | P01308 | 0 | Both | B49:01/B40:01 | 0.1983/0.5284 |
| 2 | AEDLQVGQVEL | INS | 58-68 | P01308 | 0 | Both | B40:01 | 0.1346 |
| 2 | VELGGGPGAGSL | INS | 66-77 | P01308 | 0 | Both | B40:01 | 0.327 |
| **2** | **REAEDLQGSL** | **INS-205** | **56-65** | **P01308** | **0** | **IFN-α** | **B40:01/B49:01** | **0.043/0.347** |
| 4 | AEIPGGPEA | PCSK1 | 37-45 | P29120 | 0 | IFN-α | B49:01/B40:01 | 0.0656/0.321 |
| 2 | GEDARGTWTL | PCSK2 | 561-570 | P16519 | 0 | IFN-α | B40:01 | 0.0816 |
| 4 | AERPLNEQI | SCG3 | 38-46 | Q8WXD2 | 0 | IFN-α | B49:01/B40:01 | 0.0194/0.134 |
| 4 | AEDIVHKI | SCG3 | 148-155 | Q8WXD2 | 0 | IFN-α | B49:01/B40:01 | 0.032/0.416 |
| 4 | KEKETLITI | SCG3 | 294-302 | Q8WXD2 | 0 | IFN-α | B49:01/B40:01 | 0.0067/0.100 |
| 3 | SEADIQRLL | SCG5 | 36-44 | P05408 | 0 | IFN-α | B40:01/B49:01 | 0.0104/0.0188 |
| 3 | VEYPAHQAM | SCG5 | 58-66 | P05408 | 0 | IFN-α | B40:01/B49:01 | 0.0341/0.0761 |
| 3 | AEFSREFQL | SCG5 | 138-146 | P05408 | 0 | IFN-α | B40:01/B49:01 | 0.0156/0.0449 |
| 3 | REFQLHQHL | SCG5 | 142-150 | P05408 | 0 | IFN-α | B40:01 | 0.008 |

**Supplementary Table 3. HLA-B40-restricted candidate epitopes used for screening the reactivity of TCRs and T-cell lines from the islet-infiltrating T-cells of HLA-B40$^{+}$ nPOD T1D organ donors.** TCR reactivity was screened with individual peptides, while T-cell lines where screened using the indicated peptide pools. Peptides testing positive after deconvolution of individual reactivities are marked in bold.

| nPOD case / RRID:SAMN# | Negative control peptides |
|---|---|
| 6480 / 15879533 | **Irrelevant allele: HLA-A*02:01**<br>$GAD_{114-123}$: VMNILLQYVV<br>IA-$2_{797-805}$: MVWESGCTV<br>$IGRP_{265-273}$: VLFGLGFAI<br>$PPI_{15-24}$: ALWGPDPAAA<br>$ZnT8_{186-194}$: VAANIVLTV<br>INS $DRiP_{1-9}$: MLYQHLLPL |
| 6536 / 18242780 | **Irrelevant allele: HLA-B*07:02**<br>$GAD_{3-11}$: SPGSGFWSF<br>$GAD_{100-108}$: ACDGERPTL<br>$GAD_{175-184}$: HPRYFNQLST<br>$GAD_{311-320}$: IPSDLERRIL<br>$GAD_{498-506}$: KPQHTNVCF<br>$GAD_{530-538}$: APVIKARMM<br>$PPI_{8-16}$: LPLLALLAL |
| 6563 / 30386851 | **Irrelevant allele: HLA-B*39:06**<br>$PPI_{5-12}$: MRLLPLLA<br>**Irrelevant HLA-A*02:01-binding peptide**<br>West Nile Virus $PP_{430-438}$: SVGGVFTSV |
| 6566 / 33284286 | **Irrelevant allele: HLA-B*39:06**<br>$PPI_{5-12}$: MRLLPLLA |

**Supplementary Table 4. Negative control peptide pools and individual peptides used for screening the reactivity of T-cell lines from the islet-infiltrating T-cells of HLA-B40$^+$ nPOD T1D organ donors.** Peptide(s) binding to irrelevant HLA-I alleles (i.e. not expressed by the corresponding donor) or an irrelevant viral peptide binding to the HLA-A*02:01 allele expressed by donor 6563 were used.

| **Gene** | **Forward primer** | **Reverse primer** |
|---|---|---|
| *INS* | TTCTACACACCCAAGACCCG | ATTGTTCCACAATGCCACGC |
| *INS* | TGTCCTTCTGCCATGGCCCT | TTCACAAAGGCTGCGGCTGG |
| *CHGA* | CTGAACACAGGCAGCTTTCTA | CAGTCAGGAGTTCTCAGCTTTC |
| *PCKS1* | CATTCTTTGCCTGGTGCACTGTGT | TTGTGGCTGAGAAAGGAGACAGGT |
| *PCSK2* | AACCGTGCCTGAGAGATTCC | TCGGTTGTGAGTGTCAGCAC |
| *SYT4* | CACCAGCCGGGAAGAATTTG | GAAGACCAGGCCAAATGCAC |
| *FTO* | GCATGGCTGCTTATTTCGGG | GGATGCGAGATACCGGAGTG |
| *SOX9* | TTCACCTACATGAACCCCGC | AAGGTCGAGTGAGCTGTGTG |
| *PMSB5* | GCCTTCAAGTTCCGCCAT | TGCCTAGCAGGTATGGGTTG |
| *PMSB6* | CAACCACTGGGTCCTACATCG | GGAAACCGAGCTGGTAGGTG |
| *PMSB7* | AAGTTGCCTTATGTCACCATGG | ATGGCTTCGCTCACCAGAT |
| *PMSB8* | GGCTGTACTATCTGCGAAATGG | AGTCCAGGACCCTTCTTATCCC |
| *PMSB9* | ATGCTGACTCGACAGCCTTT | GAGCAATAGCGTCTGTGGTG |
| *PMSB10* | ACTGCCAAAGAAATGCATCAT | ATCGTTAGTGGCTCGCGTAT |
| *GAPDH* | TCGTGGAAGGACTCATGACC | ATGATGTTCTGGAGAGCCCC |
| *ACTB* | ACTCTTCCAGCCTTCCTTCC | CGTACAGGTCTTTGCGGATG |
| *PPIA* | ATGGCAAATGCTGGACCCAACA | ACATGCTTGCCATCCAACCACT |

**Supplementary Table 5. Primers used to quantify gene expression of β-cell identity markers and constitutive proteasome/immunoproteasome subunits.**

| Antibody [Clone] | Supplier, Cat# | RRID | Species, Clonality | Staining conditions | Secondary Ab fluorophore | OPAL conditions |
|---|---|---|---|---|---|---|
| **HLA-B [HC10]** | Origene, AM33035PU-N | AB_2728622 | Mouse, mAb | 1:700 1 h RT | OPAL 520 | 1:100 8 min |
| **HLA-A/B/C/E [EMR8-5]** | Abcam, ab70328 | AB_1269092 | Mouse, mAb | 1:700 1 h RT | OPAL 570 | 1:80 10 min |
| **INS [ICTABLS]** | ThermoFisher, 14-9769-82 | AB_2573014 | Mouse, mAb | 1:1,500 1 h RT | OPAL 690 | 1:100 10 min |
| **HLA-A [ARC0588]** | Invitrogen, MA5-35106 | AB_2849011 | Rabbit, mAb | 1:500 1 h RT | OPAL 620 | 1:100 12 min |
| **GCG [K79bB10]** | Abcam, ab10988 | AB_297642 | Mouse, mAb | 1:800 0.5 h RT | OPAL 780 | 1:80 10 min |

**Supplementary Table 6. Abs and conditions used for multiparameter immunofluorescence staining of human pancreas tissues.** RT, room temperature.

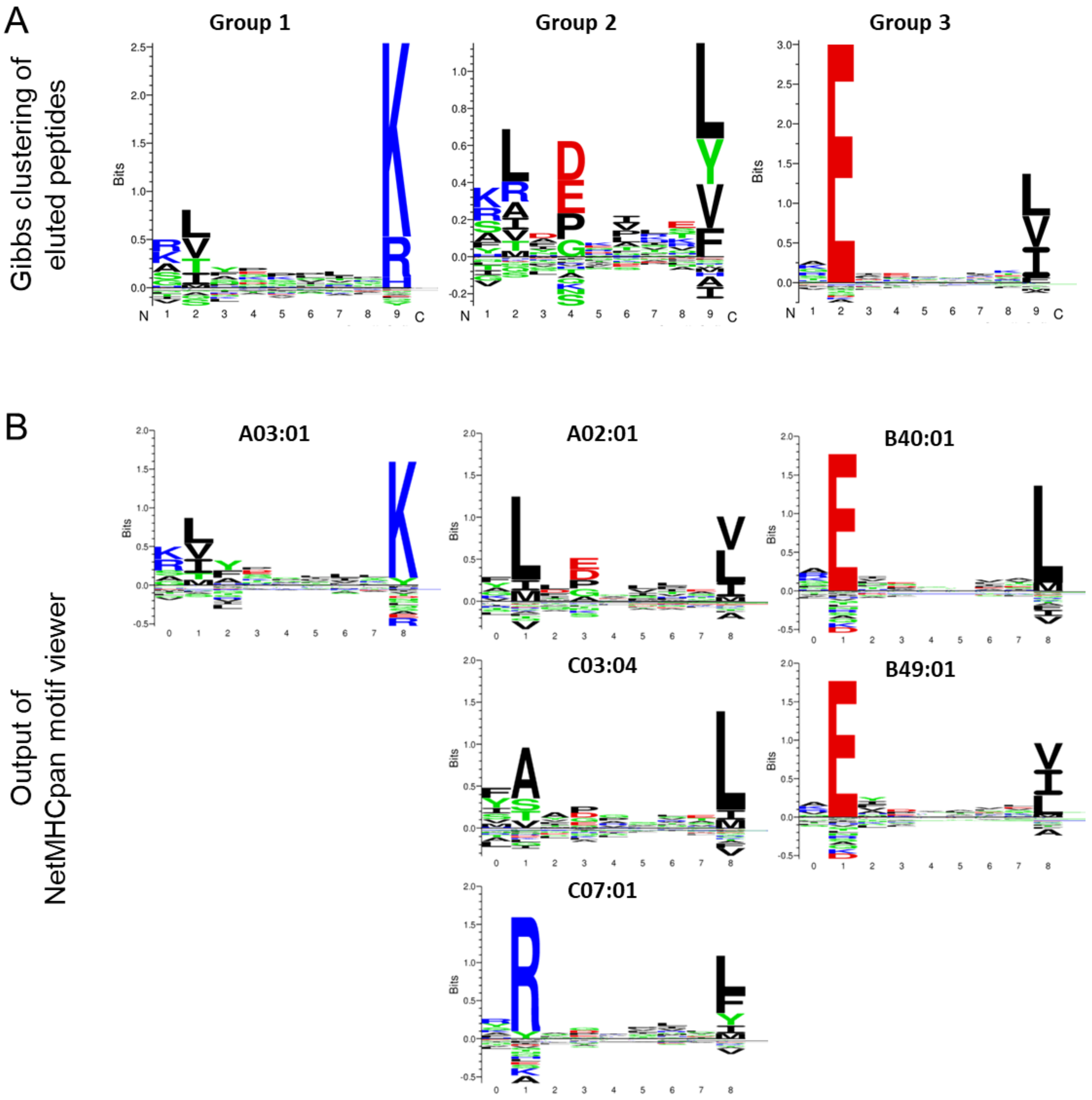


**Supplementary Figure 1. Gibbs clustering of 8-14mer peptides. A.** All unique conventional and post-translationally modified candidates identified in the 8-14 aa length range were clustered using GibbsCluster. The *x*-axis indicates the residue position within the 9mer core sequence. Each aa is represented by its single-letter code, with its size proportional to its frequency at the indicated position. All peptides were inputed as unmodified sequences. **B.** NetMHCpan4.1a sequence motifs for the HLA-I alleles expressed by the ECN90 β-cells.

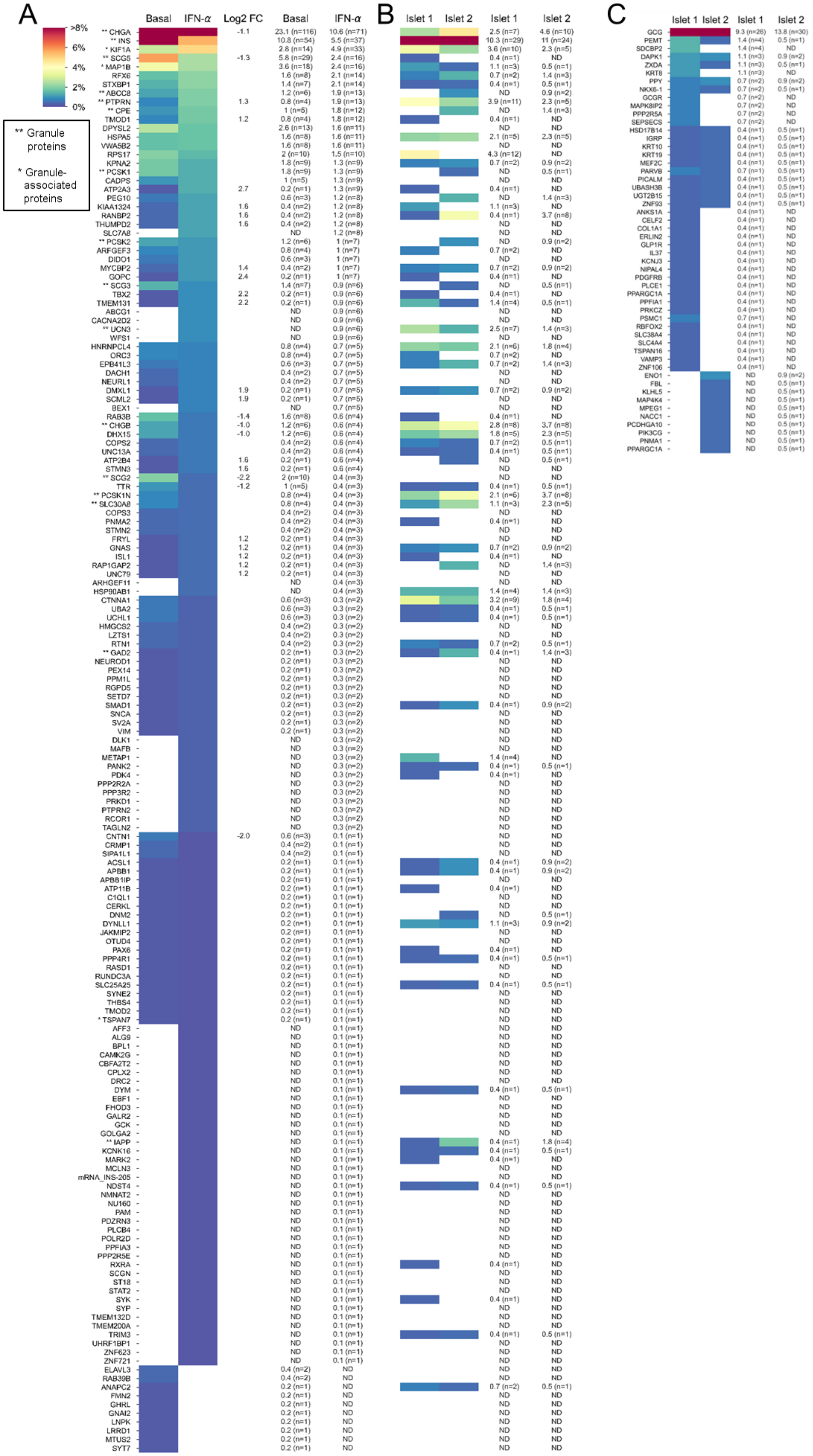

A
Basal
IFN-α
Log2 FC
Basal
IFN-α
>8%
6%
4%
2%
0%
** Granule proteins
* Granule-associated proteins
B
Islet 1
Islet 2
Islet 1
Islet 2
C
Islet 1
Islet 2
Islet 1
Islet 2

**Supplementary Figure 2. Heatmap of the source proteins of the immunopeptidome of ECN90 β-cells and primary human islets. A-B.** Relative representation of source proteins in ECN90 β-cells (A) and human islets (B), along with the corresponding significant log2 fold changes (FC) and percent values, ranked according to the number of peptides detected in the IFN-α-treated condition. **C.** Relative representation of GCG and β-cell-enriched source proteins in two different primary islet preparations. In all panels, relative representation of each source protein is calculated based on the number of unique peptides identified for each protein out of the total number of peptides in a given condition, expressed in percentage. Ligands included herein are conventional, post-translationally modified and mRNA splice peptides. Peptides carrying PTMs were counted only for PTMs defined as likely biological; they were otherwise counted as unmodified. mRNA splice variants were considered only when validated by spectral matching.

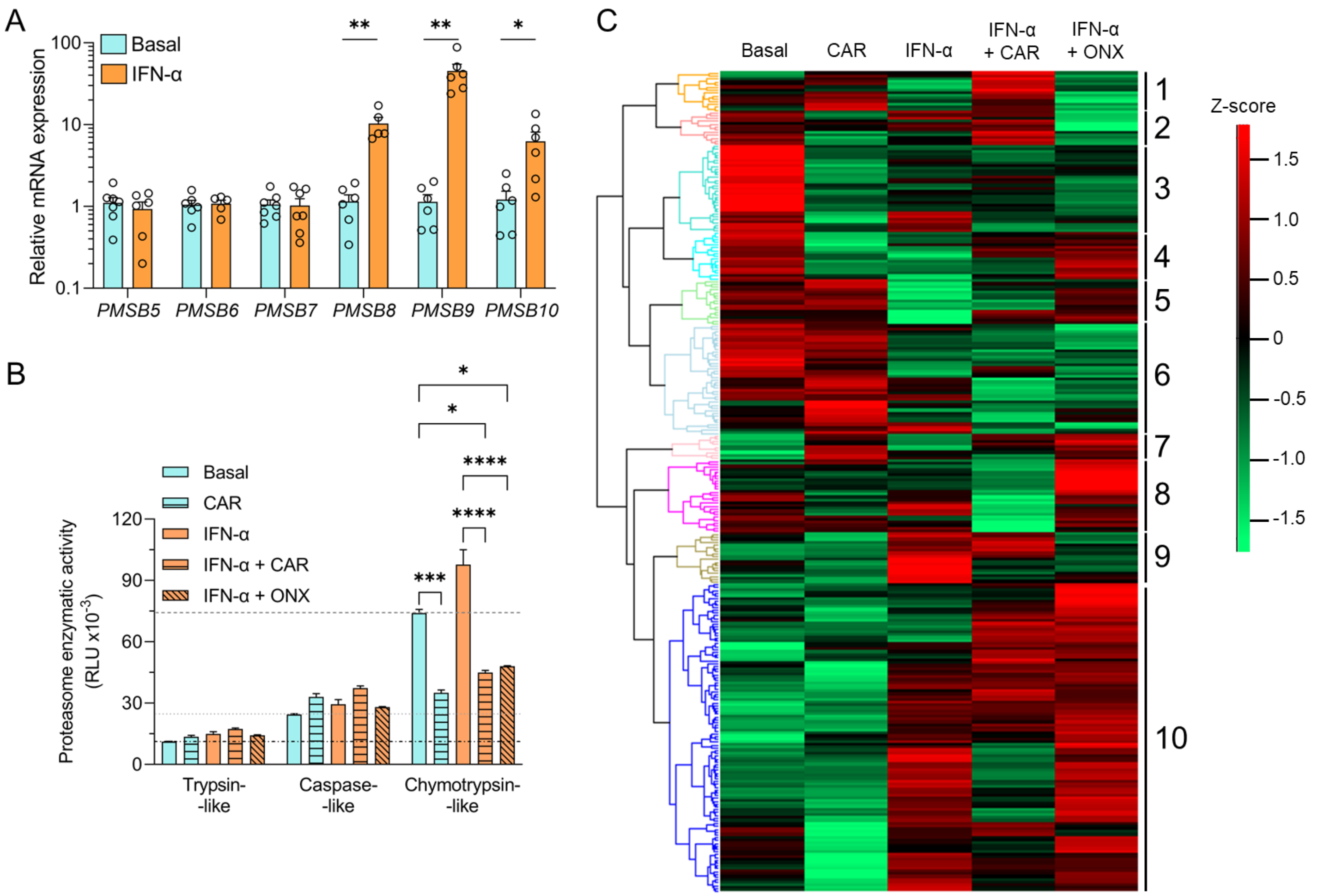


**Supplementary Figure 3. Inhibition of constitutive and immuno-proteasome in ECN90 β-cells. A.** Gene expression of the subunits of the constitutive (left; *PSMB5*, *PSMB6*, *PSMB7*) and immuno-proteasome (right; *PSMB8*, *PSMB9*, *PSMB10*) in ECN90 β-cells assessed by qPCR (n=6 biological replicates). **B.** Enzymatic activity of the three proteasome catalytic subunits in basal and IFN-α-treated ECN90 β-cells (3-5×$10^6$, n=3 biological replicates) with or without carfilzomib (CAR; constitutive proteasome inhibitor) or ONX-0914 (ONX; immuno-proteasome inhibitor). Data represent mean±SEM of triplicate measurements, with horizontal lines indicating the basal enzymatic activity of each subunit. **p*<0.01, ****p*<0.0005, *****p*<0.0001 by two-way ANOVA. **C.** Heatmap of HLA-I-eluted peptides identified in the indicated conditions. For each precursor found in ≥2 replicates in one of the 5 conditions, ion intensities were summed. For peptides identified in multiple instances, median values were computed. The values were log2-transformed, missing values were imputed from normal distribution downshifted by 1.8 and normalization was computed using row z-scores. The heatmap was generated with Perseus (v2.0.6), clustering rows with "average" as the agglomeration method and "Pearson correlation" as the distance matrix. Dendrogram colors on the left and numbers on the right indicate distinct peptide clusters (see Results). The identity of peptides in each cluster is provided in Supplementary Data 8.

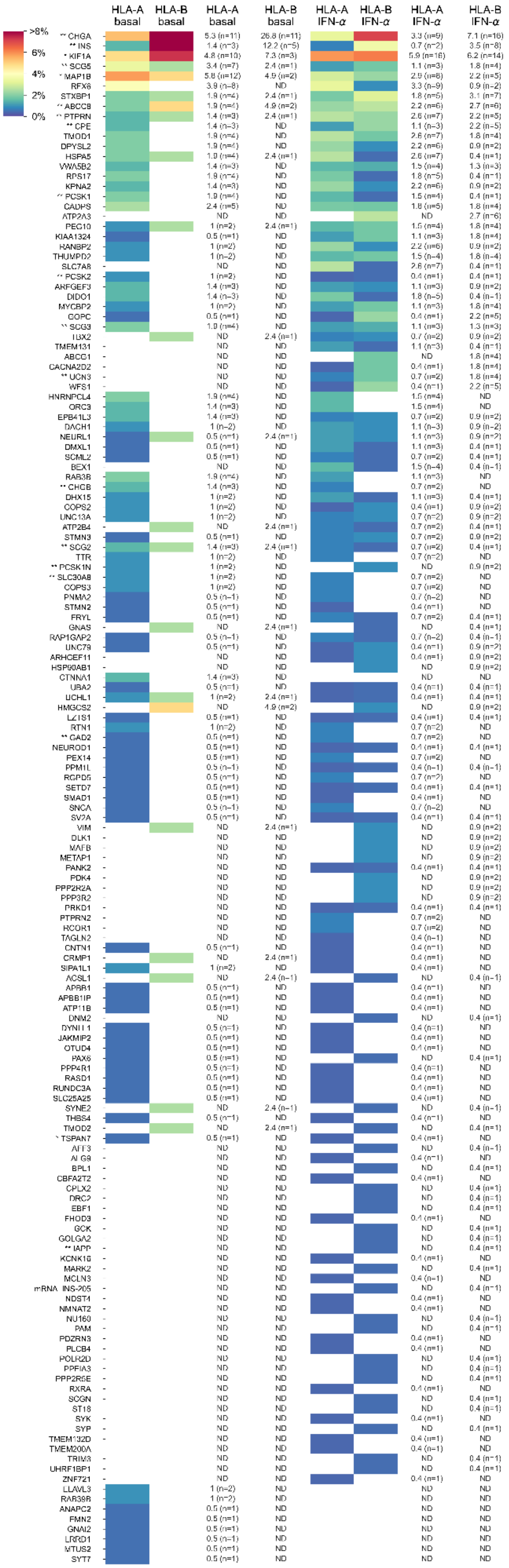


**Supplementary Figure 4.** Heatmap of the source proteins of the immunopeptidome of ECN90 β-cells separated by predicted HLA-A and HLA-B restriction. Ranking and legend is the same as in Supplementary Fig. 2.

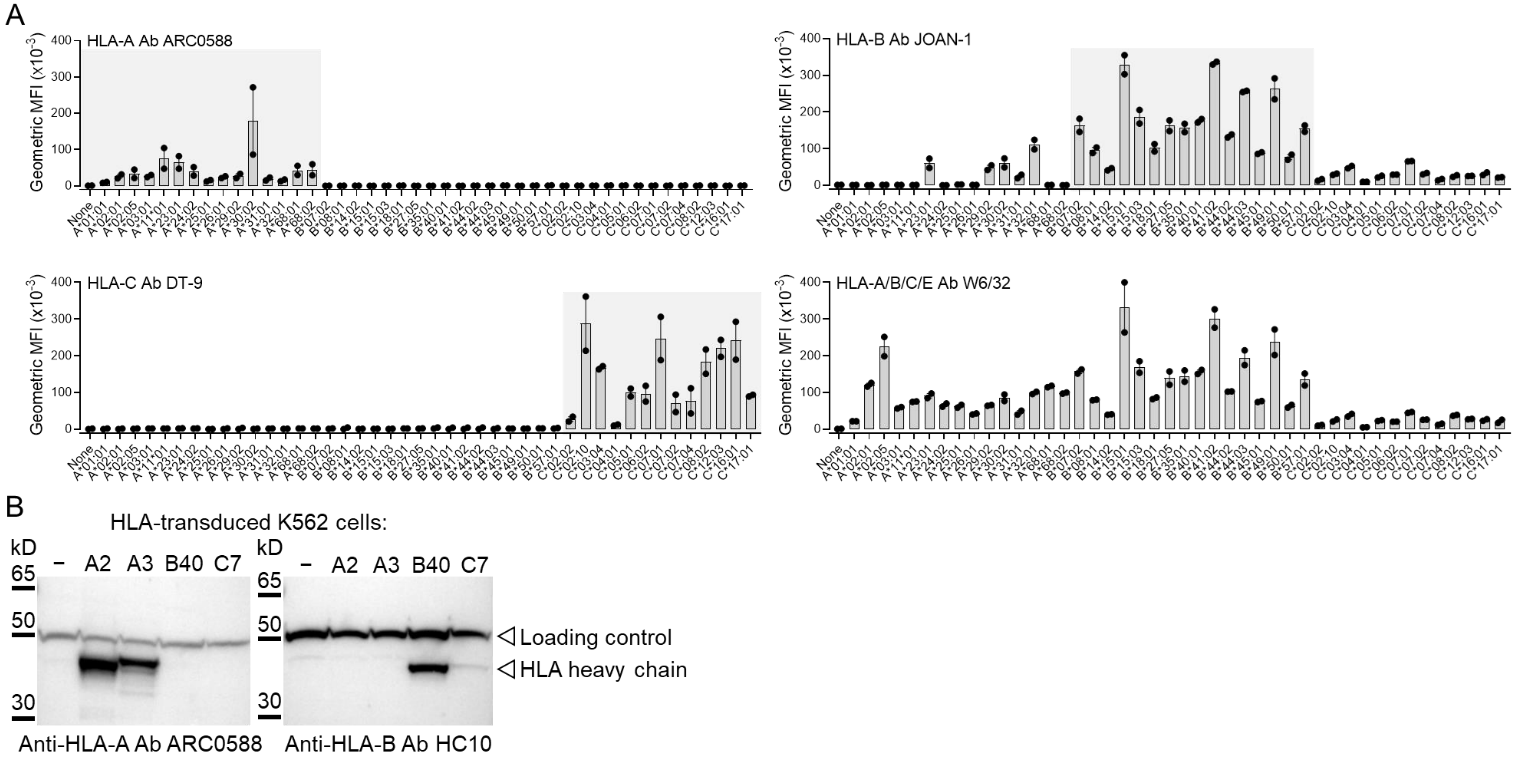


**Supplementary Figure 5. Validation of the specificity of Abs recognizing HLA-A, -B or -C. A.** Flow cytometry staining of HLA-I$^{-}$ K562 cells transduced with the indicated HLA-I alleles. Data is represented as mean±SEM of duplicate measurements from a representative experiment out of 2 performed. **B.** Western blot detection of HLA-I heavy chains with the indicated Abs on HLA-I$^{-}$ K562 cells transduced with the indicated HLA-I alleles. α-tubulin was used as loading control. A representative experiment out of 3 performed is shown.

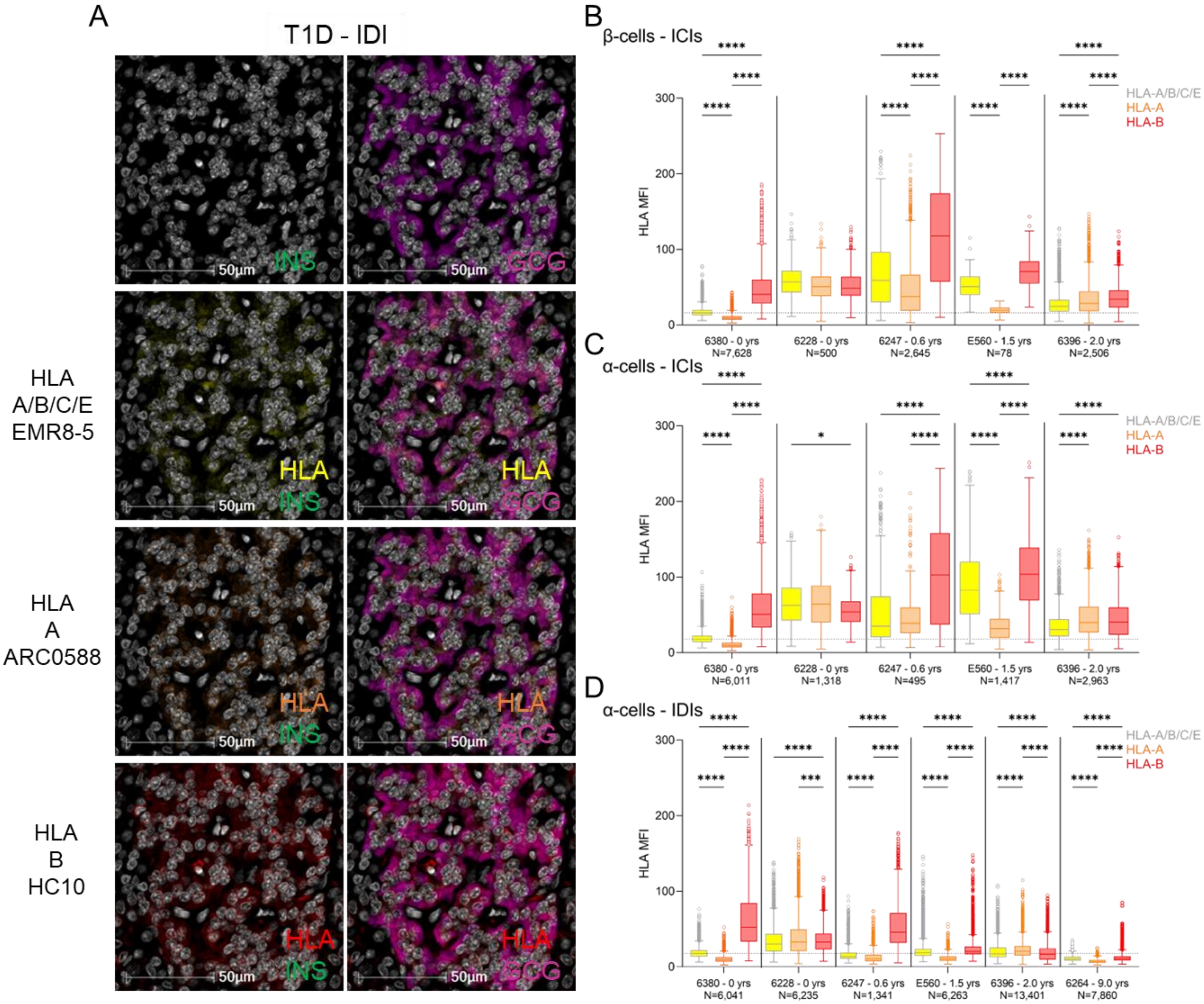


**Supplementary Figure 6. HLA-B vs. HLA-A hyper-expression in the islets of individual T1D cases. A.** Representative immunofluorescence images of IDIs from T1D case nPOD 6396, stained for INS (green, first column) or GCG (violet, second column), alone (first raw) or in combination with HLA-A/B/C/E (yellow; second raw), HLA-A (orange; third raw) and HLA-B (red, fourth raw). Scale bar 50 µm. **B-C-D.** Immunofluorescence quantification of HLA-I mean fluorescence intensity (MFI) for HLA-A/B/C/E (yellow/grey), HLA-A (orange) and HLA-B (red) in β-cells (B), ICI α-cells (C) and IDI α-cells (D) from individual T1D cases (listed in Supplementary Table 1). T1D duration and number of cells analyzed are indicated for each donor. Boxes depict median and interquartile range values, with whiskers and outliers plotted with the Tukey method. Case nPOD 6264 did not display any ICI and was therefore analyzed only for IDIs. **p*<0.05, ****p*=0.0002 and *****p*<0.0001 by Dunn's multiple comparison test.

```
MKSIYFVAGL FVMLVQGSWQ RSLQDTEEKS RSFSASQADP LSDPDQMNED KRHSQGTFTS DYSKYLDSRR AQDFVQWLMN
MKSIYFVAGL            RSLQDTEEKS R                       HSQGTFTS D          AQDFVQWLMN ▶
 KSIYFVAGL            RSLQDTEEKS RSF                     HSQGTFTS DY           DFVQWLMN ▶
                       SLQDTEEKS                         HSQGTFTS DYSK          FVQWLMN ▶
                       SLQDTEEKS RS                         GTFTS DYSK
                       SLQDTEEKS RSFS                         FTS DYSKYL
                        LQDTEEKS RSFS                          TS DYSKYL
                          DTEEKS RSFS                          TS DYSKYLDS
                          DTEEKS RSFSA                          S DYSKYLD
                          DTEEKS RSFSAS                         S DYSKYLDS
                           TEEKS RSFSA                            DYSKYLDS
                                  SFSASQAD                         YSKYLDSRR AQD
                                   FSASQADP L
                                          P LSDPDQMNE
```

```
  TKRNRNNIAK RHDEFERHAE GTFTSDVSSY LEGQAAKEFI AWLVKGRGRR DFPEEVAIVE ELGRRHADGS FSDEMNTILD
◀ TKRN        HDEFERHAE G              AAKEFI AWL
◀ T           HDEFERHAE GT                EFI AWLVKG
◀ T              FERHAE GTFT               FI AWLVKG
                 FERHAE GTFTSD
                 FERHAE GTFTSDVS
                     AE GTFTSDV
                          FTSDVSSY L
```

```
NLAARDFINW LIQTKITDRK
      FINW LIQTKITD
       INW LIQTKITD
           LIQTKITD
```

Signal peptide
GRPP
IP-1
Major proGCG fragment

**Supplementary Figure 7. Mapping of the GCG peptides eluted from HLA-I molecules of primary islet samples.**
